# Human cytomegalovirus deploys molecular mimicry to recruit VPS4A to sites of virus assembly

**DOI:** 10.1101/2024.01.04.572781

**Authors:** Benjamin G. Butt, Daniela Fischer, Alison R. Rep, Martin Schauflinger, Clarissa Read, Thomas Böck, Manuel Hirner, Stephen C. Graham, Jens von Einem

**Author notes:** Correspondence: Stephen C Graham; Jens von Einem. Contributed equally to the work (Benjamin G. Butt, Daniela Fischer, Alison R. Rep). Conceptualisation: BGB, DF, SCG, JvE; Data Curation: SCG; Funding Acquisition: SCG, JvE; Investigation: ARR, DF, BGB, MS, CR, TB, MH, SCG, JvE; Project Administration: SCG, JvE; Supervision: SCG, JvE; Visualisation: BGB, SCG, JvE; Writing – Original Draft Preparation: SCG, JvE; Writing – Review & Editing: BGB, DF, ARR, MS, CR, SCG, JvE.

## Abstract

The AAA-type ATPase VPS4 is recruited by proteins of the endosomal sorting complex required for transport III (ESCRT-III) to catalyse membrane constriction and membrane fission. VPS4A accumulates at the cytoplasmic viral assembly complex (cVAC) of cells infected with human cytomegalovirus (HCMV), the site where nascent virus particles obtain their membrane envelope. Here we show that VPS4A is recruited to the cVAC via interaction with pUL71. Sequence analysis, deep-learning structure prediction, molecular dynamics and mutagenic analysis identify a short peptide motif in the C-terminal region of pUL71 that is necessary and sufficient for the interaction with VPS4A. This motif is predicted to bind the same groove of the N-terminal VPS4A Microtubule-Interacting and Trafficking (MIT) domain as the Type 2 MIT-Interacting Motif (MIM2) of cellular ESCRT-III components, and this viral MIM2-like motif (vMIM2) is conserved across β-herpesvirus pUL71 homologues. However, recruitment of VPS4A by pUL71 is dispensable for HCMV morphogenesis or replication and the function of the conserved vMIM2 during infection remains enigmatic. VPS4-recruitment via a vMIM2 represents a previously unknown mechanism of molecular mimicry in viruses, extending previous observations that herpesviruses encode proteins with structural and functional homology to cellular ESCRT-III components.

## Introduction

Enveloped viruses have developed several mechanisms to modulate host cell membranes to support the generation of viral progeny. Many of these viruses, including herpesviruses, exploit the cellular Endosomal Sorting Complex Required for Transport (ESCRT) membrane remodelling machinery by recruiting ESCRT components to sites of viral budding (reviewed in (Rivera-Cuevas and Carruthers, 2023)). The ESCRT machinery consists of four multi-protein complexes (ESCRT-0, -I, -II and -III) plus Bro-domain containing proteins like ALIX and the AAA-ATPases vacuolar protein sorting (VPS)4A and VPS4B; together these components support membrane deformation and scission events including nuclear envelope repair and formation of intraluminal vesicles (ILVs) at multivesicular bodies (MVBs) (Christ et al., 2017; McCullough et al., 2018). Bro-domain containing proteins and the ESCRT-0, I and II complexes recognise cargo molecules destined for envelopment and initiate the sequential assembly of the ESCRT machinery at different cellular sites, culminating in the recruitment of ESCRT-III components that act as effectors of membrane remodelling and scission. The interaction of ESCRT-III subunits with VPS4 drives membrane constriction and fission, with disassembly of ESCRT-III filaments to their monomeric subunits facilitating the recycling of ESCRT proteins for further rounds of assembly (Maity et al., 2019). VPS4 is recruited to sites of ESCRT-III activity via an interaction between the N-terminal Microtubule-Interacting and Trafficking (MIT) domain of VPS4 and MIT-Interacting Motifs (MIM)s located in the C-terminal tails of ESCRT-III proteins (Wenzel et al., 2022). There are seven distinct ways in which the different MIM motifs can bind their targets, which span a diverse set of cellular proteins including ATPases, kinases and proteases (Wenzel et al., 2022).

Many enveloped viruses appropriate the cellular ESCRT machinery to promote budding of nascent virus particles at the plasma membrane or into the lumen of intracellular organelles (Votteler and Sundquist, 2013), so-called ‘inside-out’ budding that represents membrane wrapping of cytoplasmic material. Most viruses accomplish this via molecular mimicry of the short peptide motifs (P[S/T]AP, PPxY and YPxL, where x is any amino acid) used by cellular ESCRT components to promote condensation of the ESCRT machinery onto a target membrane. These short motifs are termed ‘late domains’ in the context of viral assembly given their function at late stages of virion morphogenesis (Scourfield and Martin-Serrano, 2017; Votteler and Sundquist, 2013). A prime example is the structural protein Gag from human immunodeficiency virus (HIV-1), which stimulates virion budding at the plasma membrane. TSG101 and ALIX are recruited to the plasma membrane by an interaction mediated by the PTAP motif in the p6 region of Gag (Göttlinger et al., 1991; Huang et al., 1995). Subsequently, ESCRT-III proteins charged multivesicular body protein (CHMP)2 and CHMP4 are recruited together with VPS4 to facilitate scission of the budding particle from the plasma membrane (Morita et al., 2011). Large DNA viruses such as herpesviruses also appear to rely on ESCRT functions for efficient generation of infectious virus particles. Herpesvirus particle assembly is a complex process: DNA-filled capsids formed in the nucleus associate with the inner nuclear membrane and bud into the perinuclear space (so-called ‘primary envelopment’) before fusing with the outer nuclear membrane (de-envelopment), whereupon the cytoplasmic nucleocapsid associates with structural (tegument) proteins and buds into glycoprotein-studded membranes of intracellular vesicles/organelles (so-called ‘secondary envelopment’) that eventually fuse with the cell surface to release a mature enveloped virion (Ahmad and Wilson, 2020; Close et al., 2018; Owen et al., 2015; Read et al., 2019). There is increasing evidence for the involvement of ESCRT in the nuclear egress of herpes simplex virus 1 (HSV-1, a.k.a. HHV1) and Epstein Barr virus (EBV, a.k.a. HHV4). In both cases, proteins of the viral nuclear egress complex (NEC) recruit the ESCRT protein CHMP4 and ESCRT-associated protein ALIX to the inner nuclear membrane, where the NEC mediates primary envelopment of nucleocapsids (Arii et al., 2018; Lee et al., 2012). Furthermore, knockdown of ALIX and CHMP4 in the case of HSV-1 causes accumulation of capsids in the nucleus, supporting a role for ESCRT in herpesvirus nuclear egress (Arii et al., 2018). In addition to nuclear egress, HSV-1 was shown to require VPS4 ATPase activity for the formation of infectious virus particles in the cytoplasm (Calistri et al., 2007; Crump et al., 2007). The expression of dominant-negative VPS4 leads to stalled late stages of the envelopment process, especially membrane fusion, while the initiation of secondary envelopment is not affected.

Secondary envelopment of herpesviruses at intracellular glycoprotein-studded membranes is topologically similar to the formation of ILVs in MVBs, with membrane wrapping of the tegument- decorated capsid to form a bud neck that is then resolved to yield a mature virus particle in the lumen of an intracellular compartment (Read et al., 2019). Interestingly, several herpesvirus tegument and glycoproteins contain late domains through which interaction with ESCRT is possible, e.g. the conserved large tegument protein HSV-1 pUL36 (Calistri et al., 2007). Late domains are also found in various proteins of the human β-herpesvirus 5, better known as human cytomegalovirus (HCMV) (Fraile-Ramos et al., 2007). While many of these proteins are implicated in secondary envelopment, the precise role of ESCRT in HCMV infection remains controversial. Early work with siRNA knockdown showed that ESCRT is not involved in virus morphogenesis (Fraile-Ramos et al., 2007). In contrast, expression of dominant-negative VPS4 and dominant-negative ESCRT-III protein CHMP1B resulted in severely restricted virus growth (Tandon et al., 2009). These latter results are contradicted by a more recent study using inducible expression of dominant-negative VPS4 and dominant-negative ESCRT-III proteins, which showed that production of infectious virions and HCMV envelopment do not require the ESCRT machinery (Streck et al., 2020).

Although there is increasing evidence that VPS4 and ESCRT have no dominant proviral role during HCMV infection, VPS4 and other ESCRT components such as CHMP1 and HRS localize in the vicinity of the cytoplasmic viral assembly compartment (cVAC) where HCMV viral assembly and maturation in the cytoplasm takes place (Das and Pellett, 2011; Moorman et al., 2010; Streck et al., 2020; Tandon et al., 2009). The cVAC is formed during HCMV infection at the indentation of a kidney-shaped nucleus. It consists of intertwined membranes of different origins arranged in a concentric pattern around the microtubule organizing centre (Alwine, 2012; Das et al., 2007; Das and Pellett, 2011; Homman-Loudiyi et al., 2003; Sanchez et al., 2000b, 2000a). The peripheral region of the cVAC consists of Golgi-derived membranes, whereas the central part is composed of membranes positive for markers of early and late endosomes (Das et al., 2007). The importance of the cVAC for virus assembly and maturation is emphasised by the high abundance of the components (nucleocapsids, proteins and membranes) required for virion assembly, the observation of secondary envelopment only in the region of the cVAC (Schauflinger et al., 2013), and the fact that disruption of cVAC biogenesis leads to defects in virus growth (Rebmann et al., 2016). Several viral proteins have already been found to be involved in cytoplasmic virus assembly, of which pUL71 and its interaction partner pUL103 (Butt et al., 2020) play a special role as they are particularly involved in the late stages of secondary envelopment (Ahlqvist and Mocarski, 2011; Das et al., 2014; Read et al., 2019; Schauflinger et al., 2011; Womack and Shenk, 2010). Deficiencies in these proteins result in an accumulation of nucleocapsids at various advanced stages of envelopment, which is consistent with impaired envelopment and a block in membrane scission at the end of the envelopment process. However, the precise molecular mechanism by which pUL71 remodels membranes or mediates membrane scission remains to be elucidated. The ability of pUL71 to assemble into oligomers or protein complexes (Meissner et al., 2012), the membrane association of pUL71, and its endocytic trafficking appear to be important factors for its function in secondary envelopment, with the latter being important for pUL71 localisation at the cVAC (Dietz et al., 2018). Interestingly, pUL71 and pUL103 are conserved among herpesviruses and the homologues share similar functions (Fuchs et al., 2005; Jiang et al., 2017; Klupp et al., 2005). For example, the HSV- 1 tegument proteins pUL51 (pUL71 homologue) and pUL7 (pUL103 homologue) form a complex and promote virus assembly by stimulating cytoplasmic envelopment of capsids (Albecka et al., 2017; Nozawa et al., 2005; Roller and Fetters, 2015). These similarities imply that herpesviruses have evolved conserved mechanisms for important steps in virion morphogenesis, including the secondary envelopment process. This is further supported by the very similar crystal structures of the homologous complexes pUL7:pUL51 from HSV-1 (Butt et al., 2020) and BBRF2:BSRF1 from EBV (He et al., 2020). In addition, the N-terminal domain of HSV-1 pUL51 shows striking structural homology to the α-helical N-terminal structure of the cellular ESCRT-III protein CHMP4B, and pUL51 polymerises in a CHMP4B-like manner (Butt et al., 2020). The N-terminal region of HCMV pUL71 is predicted to adopt an α-helical fold similar to HSV-1 pUL51 (Bogdanow et al., 2023). The structural similarity between the N termini of pUL51 and ESCRT-III proteins implies that pUL51 and homologues may act as viral ESCRT-III components, consistent with their role in secondary envelopment.

The C-terminal regions of pUL51 and homologues are known or predicted to be intrinsically unstructured (Butt et al., 2020) and are functionally uncharacterised, whereas the C-terminal regions of ESCRT-III proteins are known to contain MIM motifs that bind the MIT domain of VPS4 to recruit ATPase activity to sites of membrane remodelling. It had previously been hypothesised that HCMV tegument protein pUL71 may be involved in recruiting VPS4 to the cVAC (Streck et al., 2018). Here we identify a short linear motif in the C-terminal region of pUL71 with striking resemblance to the cellular Type 2 MIM (MIM2) consensus sequence present in ESCRT-III proteins including CHMP4B and CHMP6 (Kieffer et al., 2008). We show that residues 300–325 of pUL71 bind directly to the MIT domain of VPS4A, we combine structurally-informed mutagenesis with interaction and localisation studies to confirm that the pUL71 MIM2-like motif is necessary and sufficient for recruitment of VPS4A to the HCMV cVAC, and we demonstrate that this interaction is conserved in β-herpesviruses but not α- or γ-herpesviruses. Our study identifies a novel and previously unknown mechanism of viral interaction with the ESCRT machinery, extending the identification of HSV-1 pUL51 as a viral ESCRT-III like protein (Butt et al., 2020) by showing that the equivalent protein in HCMV has even closer functional homology to cellular ESCRT-III components.

## Results

### HCMV pUL71 interacts with the MIT domain of VPS4A

The N-terminal domain of pUL71, spanning residues 23 to 172, is highly conserved between β- herpesviruses (Fig. 1A). This region is predicted to be comprised primarily of α-helices (Butt et al., 2020) and largely corresponds to the CHMP-like structured regions of HSV-1 pUL51 (Butt et al., 2020) and EBV BSRF1 (He et al., 2020). The C-terminal region of pUL71 is predicted to be unstructured and is poorly conserved amongst β-herpesviruses. However, close inspection of the pUL71 amino acid sequence revealed a motif spanning residues 312–323 with close resemblance to the MIM2 sequence of cellular VPS4-interacting proteins like CHMP6 (Fig. 1A). Co-transfection experiments demonstrated that pUL71 recruits FLAG-tagged VPS4A to a juxtanuclear membranous compartment pUL71 resides, previously identified to be *trans*-Golgi (Fig. 1B) (Butt et al., 2020; Dietz et al., 2018), whereas VPS4A- FLAG retains its diffuse cytoplasmic distribution when co-transfected with a GFP-tagged form of the ERGIC-resident HCMV protein pp28 (Sanchez et al., 2000b). The human CHMP6 MIM2 adopts an extended conformation that extends along the length of the VPS4 MIT domain (Fig. 1C) and mutation of the key hydrophobic residue valine 173 to aspartic acid, or prolines 171 and 174 to alanine, are sufficient to disrupt this interaction (Kieffer et al., 2008). HCMV pUL71 mutants P315A+P318A (PPAA) and V317D retain their characteristic Golgi-like localisation but lose the ability to recruit VPS4A-FLAG to these membranes (Fig. 1D). Similarly, immunoprecipitation experiments show VPS4A-FLAG to be robustly co-precipitated by wild-type pUL71 but not by the PPAA and V317D mutants (Fig. 1E). Taken together, these results are consistent with pUL71 binding VPS4A via an interaction between the VPS4A MIT domain and the potential pUL71 MIM2 region.

**Figure 1.**
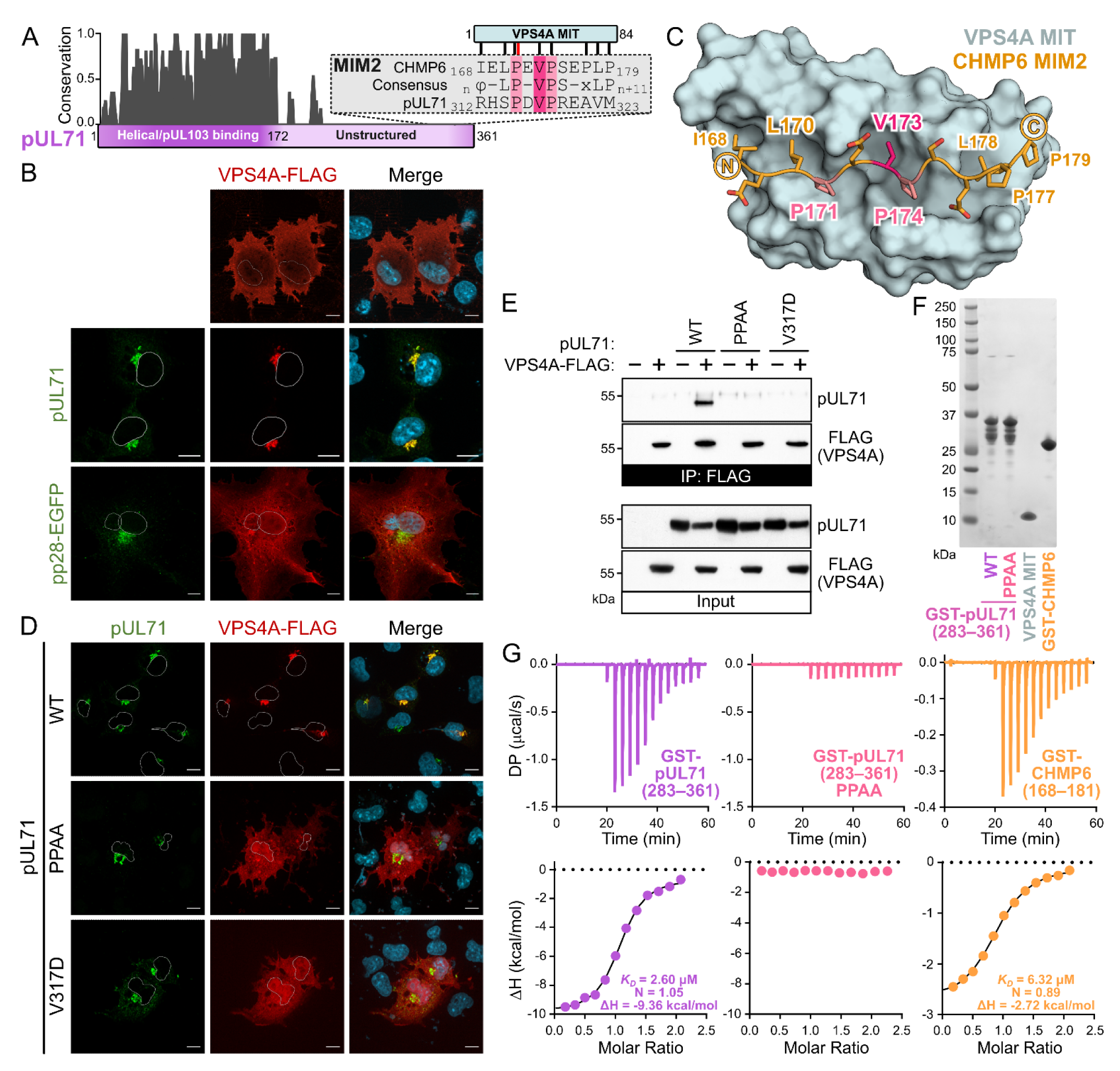
HCMV pUL71 binds the VPS4A MIT domain via a potential MIM2 motif. (A) Normalised sequence conservation of pUL71 homologues across *Betaherpesvirinae*. Inset shows an alignment of the canonical MIM2 sequence of human CHMP6, the potential MIM2 of pUL71, and the MIM2 consensus sequence where x denotes any residue, – an acidic residue and ϕ a hydrophobic residue (Kieffer et al., 2008). Vertical lines denote hydrophobic (black) and backbone hydrogen bond (red) interactions between residues of the CHMP6 MIM2 and the MIT domain of VPS4A (Kieffer et al., 2008). Selected MIM2 residues important for the interaction with VPS4A are highlighted in pink. (B) Co- transfection of pUL71 and pp28-EGFP with VPS4A-FLAG. Nuclei are outlined in single channel images and shown (DAPI, blue) in merge. Scale bars = 10 µm. (C) Structure of human CHMP6 MIM2 (orange carbon atoms, selected key residues highlighted in pink) in complex with the MIT domain of human VPS4A (PDB ID 2K3W) (Kieffer et al., 2008). (D) Co-transfection of VPS4A-FLAG with wild-type (WT) pUL71 or two mutants, P315A+P318A (PPAA) and V317D, where key residues of the pUL71 potential MIM2 were mutated and their ability to recruit VPS4A to juxtanuclear compartments is disrupted.

To probe whether pUL71 directly binds the VPS4A MIT domain, both the VPS4A MIT domain (residues 1–84) and a fusion between GST and the C-terminal region of pUL71 (spanning residues 283–361) were purified following recombinant bacterial expression (Fig. 1F). While some degradation of GST- pUL71(283–361) is evident (Fig. 1F), presumably arising from partial proteolysis of the unstructured pUL71 tail, isothermal titration calorimetry (ITC) shows GST-pUL71(283–361) to bind the VPS4A MIT domain with 2.84 ± 0.33 µM affinity (±SD, n=2; Fig. 1G and Table 1). This is tighter than the 5.54 ± 1.10 µM (±SD, n=2) interaction measured between VPS4A MIT and a GST fusion of the CHMP6 MIM2 (residues 168–181) (Fig. 1G and Table 1), and tighter than previous measurements of the CHMP6 MIM2:VPS4A MIT interaction (Kieffer et al., 2008; Wenzel et al., 2022). Purified GST-pUL71(283–361) PPAA lacks the ability to bind the VPS4A MIT domain directly (Fig. 1G). These experiments confirm that pUL71 binds the MIT domain of VPS4A directly with high affinity, and that key residues of the potential pUL71 MIM2 are essential for this interaction.

**Table 1.**
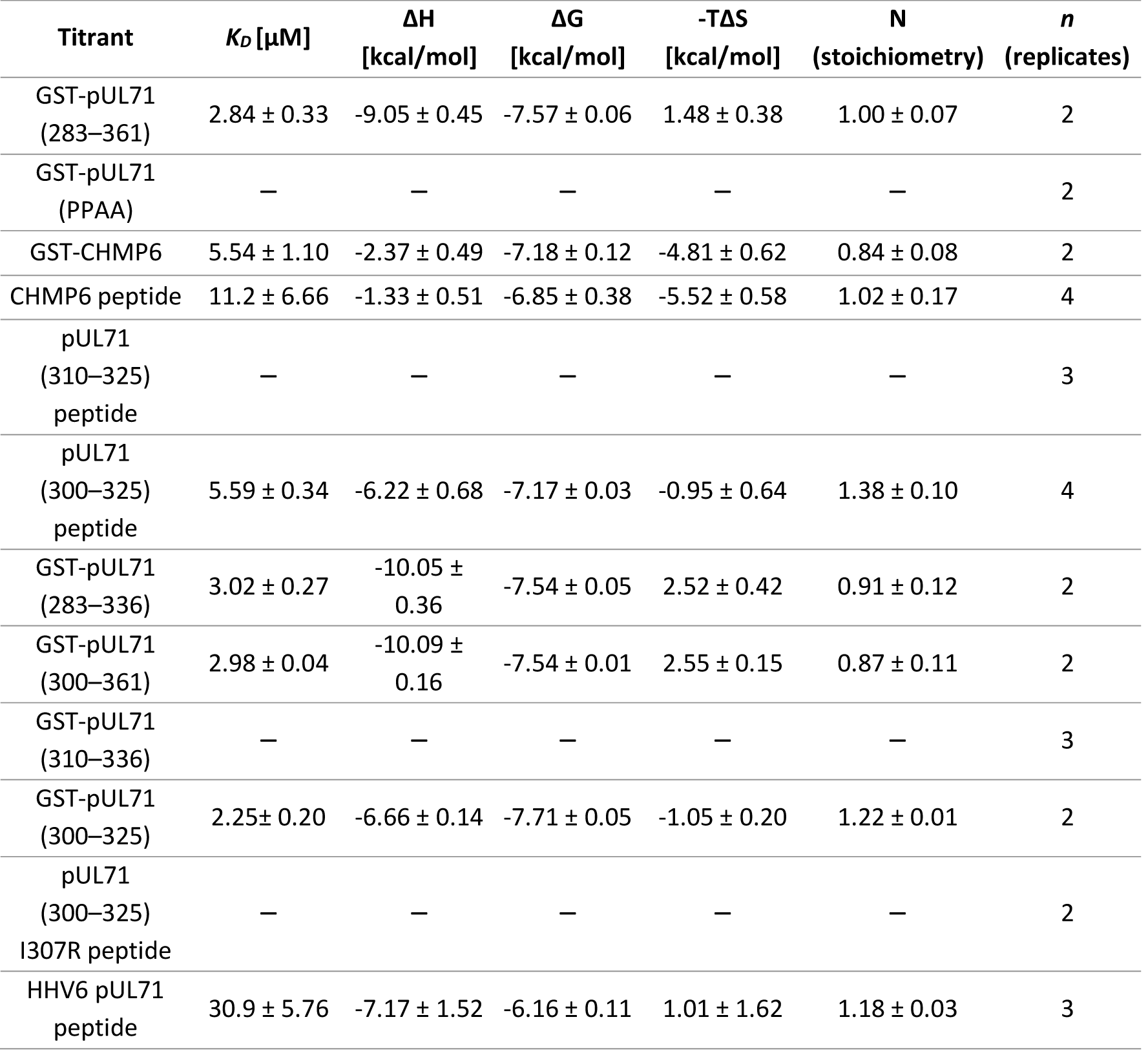
Thermodynamic properties of the interactions with VPS4 MIT domain. As quantitated by isothermal titration calorimetry (ITC). Experiments were performed *n* (replicates) times and mean ± SD values are shown. Data for individual titrations are presented as supporting information (Table S1). –, no binding detected.

| Titrant | $K_D$ [μM] | $\Delta H$<br>[kcal/mol] | $\Delta G$<br>[kcal/mol] | $-T\Delta S$<br>[kcal/mol] | N<br>(stoichiometry) | *n*<br>(replicates) |
| --- | --- | --- | --- | --- | --- | --- |
| GST-pUL71<br>(283–361) | $2.84 \pm 0.33$ | $-9.05 \pm 0.45$ | $-7.57 \pm 0.06$ | $1.48 \pm 0.38$ | $1.00 \pm 0.07$ | 2 |
| GST-pUL71<br>(PPAA) | – | – | – | – | – | 2 |
| GST-CHMP6 | $5.54 \pm 1.10$ | $-2.37 \pm 0.49$ | $-7.18 \pm 0.12$ | $-4.81 \pm 0.62$ | $0.84 \pm 0.08$ | 2 |
| CHMP6 peptide | $11.2 \pm 6.66$ | $-1.33 \pm 0.51$ | $-6.85 \pm 0.38$ | $-5.52 \pm 0.58$ | $1.02 \pm 0.17$ | 4 |
| pUL71<br>(310–325)<br>peptide | – | – | – | – | – | 3 |
| pUL71<br>(300–325)<br>peptide | $5.59 \pm 0.34$ | $-6.22 \pm 0.68$ | $-7.17 \pm 0.03$ | $-0.95 \pm 0.64$ | $1.38 \pm 0.10$ | 4 |
| GST-pUL71<br>(283–336) | $3.02 \pm 0.27$ | $-10.05 \pm 0.36$ | $-7.54 \pm 0.05$ | $2.52 \pm 0.42$ | $0.91 \pm 0.12$ | 2 |
| GST-pUL71<br>(300–361) | $2.98 \pm 0.04$ | $-10.09 \pm 0.16$ | $-7.54 \pm 0.01$ | $2.55 \pm 0.15$ | $0.87 \pm 0.11$ | 2 |
| GST-pUL71<br>(310–336) | – | – | – | – | – | 3 |
| GST-pUL71<br>(300–325) | $2.25 \pm 0.20$ | $-6.66 \pm 0.14$ | $-7.71 \pm 0.05$ | $-1.05 \pm 0.20$ | $1.22 \pm 0.01$ | 2 |
| pUL71<br>(300–325)<br>I307R peptide | – | – | – | – | – | 2 |
| HHV6 pUL71<br>peptide | $30.9 \pm 5.76$ | $-7.17 \pm 1.52$ | $-6.16 \pm 0.11$ | $1.01 \pm 1.62$ | $1.18 \pm 0.03$ | 3 |

Truncations of pUL71 were designed to identify the precise sequence motifs required for VPS4A binding. Immunofluorescence analysis of co-transfected cells showed that pUL71 with C-terminal truncations that included the entire potential MIM2 [pUL71(1–326)] or spanned all but the last three residues of the motif [pUL71(1–320)] retained the ability to recruit VPS4A-FLAG to juxtanuclear compartments (Fig. 2A). pUL71 where all but the first two residues of the potential MIM2 have been removed [pUL71(1–314)] was unable to recruit VPS4-FLAG, as was a construct with the majority of the potential MIM2 removed [pUL71(Δ315–326)] (Fig. 2A). These results demonstrate that residues 314– 320 are necessary for VPS4A binding. ITC analysis confirmed that a purified peptide encompassing the CHMP6 MIM2 (residues 168–181) was sufficient to bind purified VPS4A MIT domain with 11.2 ± 6.66 µM affinity (±SD, n=2; Fig. 2B and Table 1). However, a peptide spanning the potential pUL71 MIM2 (residues 310–325) failed to bind VPS4A (Fig. 2B), suggesting that the potential MIM2 is necessary but not sufficient to drive the interaction. Further ITC analysis of pUL71 truncations purified as GST fusions (Fig. S1 and Table 1) demonstrated that pUL71 residues 300–310 are necessary for the VPS4A interaction, in addition to the potential MIM2. ITC analysis confirmed that a peptide spanning pUL71 residues 300–325, encompassing both a predicted helical region and the potential MIM2, was sufficient to bind the VPS4A MIT domain with an affinity of 5.59 ± 0.34 µM (±SD, n=4; Fig. 2C,D and Table 1).

**Figure 2.**
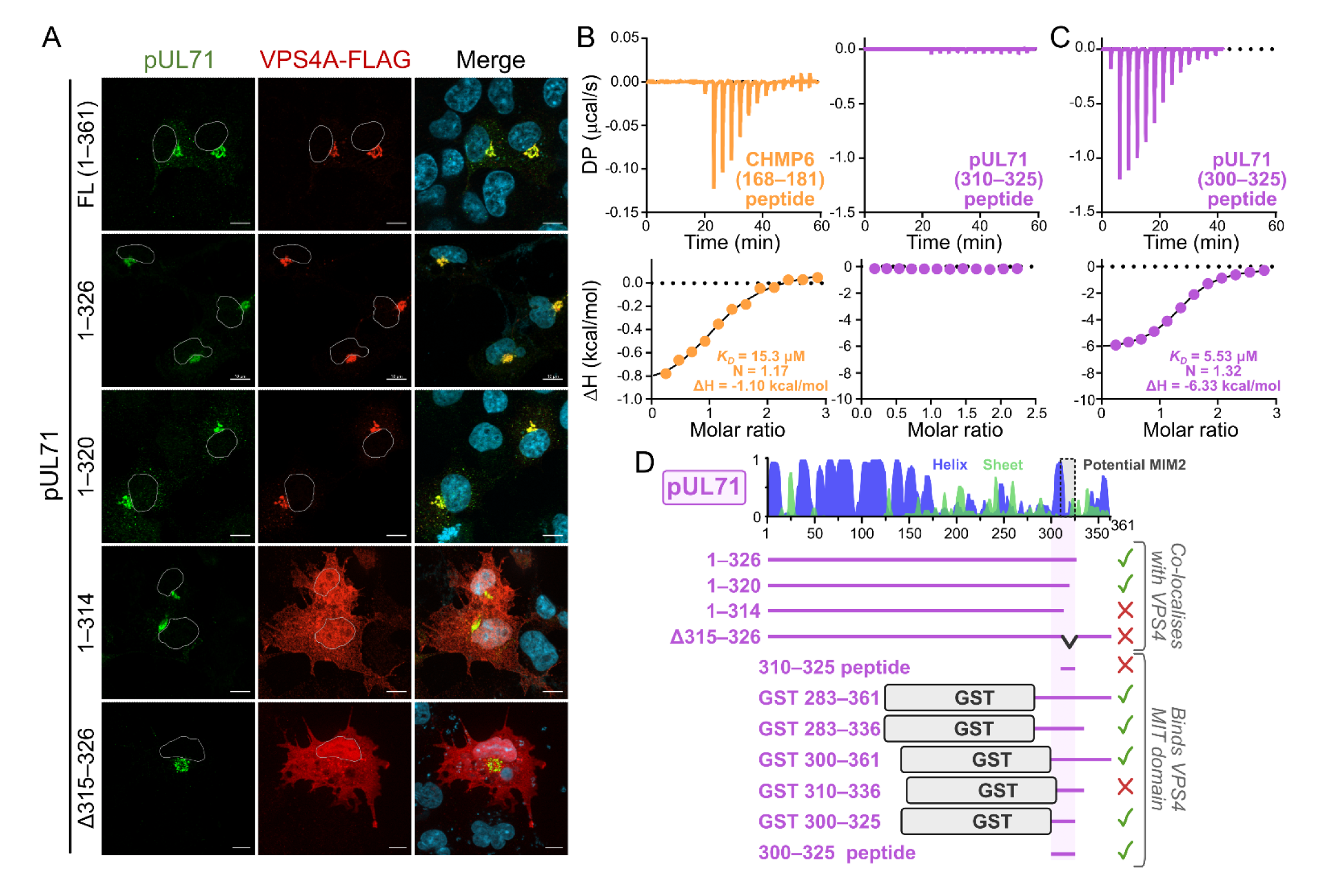
An extended MIM2-like motif spanning pUL71 residues 300–325 is necessary and sufficient for VPS4A binding. (A) Co-transfection of VPS4A-FLAG with full-length or truncated pUL71, or with pUL71 lacking the potential MIM2 (Δ315–326). Nuclei are outlined in single channel images and shown (DAPI, blue) in merge. Scale bar = 10 µm. (B) ITC analysis of the interaction between purified VPS4A MIT domain and synthetic peptides corresponding to the CHMP6 MIM2 (*left*) or the potential pUL71 MIM2 spanning residues 310–325 (*right*). (C) ITC analysis of the VPS4A MIT domain binding a peptide corresponding to the extended MIM2-like motif spanning pUL71 residues 300–325. (D) Schematic diagram of pUL71 truncation experiments. *Top:* Predicted pUL71 secondary structure is shown (blue helices and green sheets) with the potential MIM2 boxed. *Bottom:* The ability to bind VPS4A as evidenced by co-localisation following co-transfection or by ITC analysis, with residues 300–325 that are sufficient for binding highlighted (see also Fig. S1).

### Model of the HCMV pUL71 in complex with VPS4A MIT

Recent advances in deep learning have revolutionised the structural prediction of proteins and their complexes (Evans et al., 2021; Jumper et al., 2021), including the prediction of viral proteins with multiple novel domains (Gao et al., 2022) and of virus protein complexes (Benedyk et al., 2022). AlphaFold-Multimer was thus used to generate a model of pUL71(300–325) in complex with the VPS4A MIT domain (Fig. 3A). The complex is predicted with high per-residue confidence (pLDDT; Fig. 3B) and low predicted aligned error (PAE) of VPS4A residues with respect to pUL71 (Fig. 3C), consistent with a confidently predicted model of the complex. pUL71 is predicted to bind the groove between helices α1 and α3 of the VPS4A MIT domain as a short helix followed by a stretch of residues in an extended conformation. Closer inspection of the interface shows that residues I307, L308 and M311 in the short helix of pUL71 are predicted to bind a surface on VPS4A centred on residues L6 and F39 (Fig. 3A). Residues 314–322 of pUL71 adopt a similar conformation to the equivalent residues of the CHMP6 MIM2 domain, with P315, V317 and P318 predicted to lie in shallow surface pockets (Fig. 3D). PDBePISA analysis (Krissinel and Henrick, 2007) demonstrates that the predicted interaction of pUL71(300–325) with the VPS4A MIT domain has a larger interface area (912 Å^2^ versus 667 Å^2^), with more predicted hydrogen bonds (5 versus 1) and salt bridges (2 versus 1) when compared to the structure of VPS4A MIT domain with CHMP6 (Kieffer et al., 2008). This is consistent with a larger enthalpic contribution to binding (ΔH = -6.7 kcal/mol versus -2.4 kcal/mol; Fig. 2B,C and Table 1). While the MIM2 of CHMP6 binds the VPS4 MIT domain as an extended peptide (Kieffer et al., 2008), a helix- plus-extended conformation of Vps4 binding is observed for the MIM2 of Vps20 (Kojima et al., 2016), the yeast CHMP6 homologue, and for the yeast protein Vfa1 that is proposed to positively regulate Vps4 activity (Vild and Xu, 2014) (Fig. 3D). Vfa1 is reported to bind Vps4 approximately 100-fold more tightly than does Vps20 (1.8 µM vs 188 µM *K*_D_), suggesting that the presence of a helix before the extended MIM2-like region does not define interaction affinity *per se* (Kojima et al., 2016). The Type 7 MIM(N) motif of yeast Atg13 also adopts a helix-plus-extended conformation and binds between helices α1 and α3 when interacting with the second of the two tandem MIT domains of Atg1 (Fujioka et al., 2014), but the Atg13 MIM(N) and pUL71 MIM2-like motifs run in opposite directions (Fig. 3E). Mutation of VPS4A MIT residue valine 13 to aspartic acid (V13D) specifically prevents binding of MIM2 (Kieffer et al., 2008), whereas mutation of leucine 64 to aspartic acid (L64D) on the opposite face of the domain prevents binding of helical MIM1 motifs (Stuchell-Brereton et al., 2007). Co-transfection experiments confirm that the MIT domain of VPS4A is required for association with pUL71 at juxtanuclear compartments and that the V13D mutation prevents association, whereas VPS4A(L64D)- FLAG is efficiently recruited by pUL71 (Fig. 3G). Taken together, these results confirm that pUL71 binds between helices α1 and α3 of the VPS4A MIT domain.

**Figure 3.**
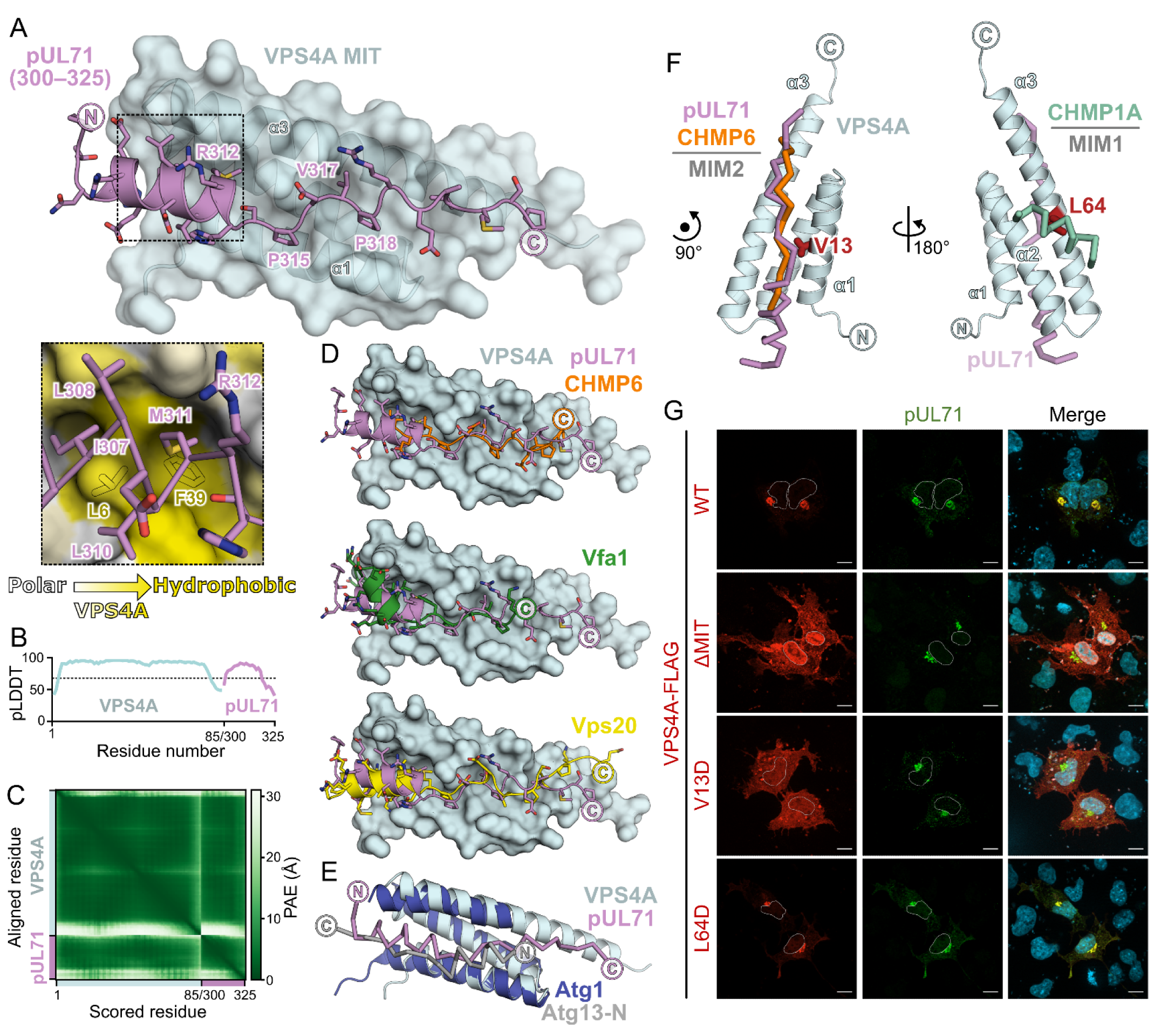
HCMV pUL71 is predicted to bind the MIM2-binding groove of the VPS4A MIT domain as a helix plus extended peptide. (A) Predicted structure of pUL71 residues 300–325 (violet ribbon with side chains shown) in complex with the VPS4A MIT domain (cyan ribbons and semi-transparent molecular surface). pUL71 is predicted to bind the groove between VPS4A helices α1 and α3. *Inset:* Predicted interaction between hydrophobic residues of the pUL71 helical region and the VPS4A MIT domain. VPS4A molecular surface is coloured by residue hydrophobicity from white (polar) to yellow (hydrophobic) and selected hydrophobic side chains are shown as silhouettes. (B) Per-residue predicted Local Distance Difference Test (pLDDT) scores for predicted complex. Values above 70 (dashed line) represent regions predicted with high confidence. (C) Predicted aligned error (PAE) matrix for predicted complex, demonstrating high confidence (low PAE, green) in relative orientation of pUL71(300–325) with respect to VPS4A MIT domain. (D) Structural comparison of the predicted pUL71(300–325) (violet ribbon and sticks) and the VPS4A MIT domain (cyan surface) complex to experimental structures of MIT domains bound to cellular MIM2s. Experimental structures were superposed by structural alignment of the MIT domains but, for clarity, only the predicted VPS4A MIT domain is shown. *Top:* Canonical CHMP6 MIM2 (orange ribbon and sticks) in complex with human VPS4A (PDB ID 2K3W) (Kieffer et al., 2008). *Middle:* Helix and MIM2 of yeast Vps4 regulator Vfa1 (green ribbon and sticks) bound to yeast Vps4 (PDB ID 4NIQ, chains A+C) (Vild and Xu, 2014). *Bottom:* Helix and non-canonical MIM2 of yeast CHMP6 homologue Vps20 (yellow ribbon and sticks) bound to yeast Vps4 (PDB ID 5FVL, chains A+D) (Kojima et al., 2016). (E) Crystal structure of the second of the two tandem MIT domains of yeast Atg1 (dark blue ribbon) in complex with the Type 7 MIM(N) of yeast Atg13 (grey C^α^ trace) (PDB ID 4P1N) (Fujioka et al., 2014) superposed on the predicted structure of human VPS4A MIT domain (cyan ribbons) and pUL71 (violet C^α^ trace). While both MIM-like motifs bind the groove between MIT domain helixes α1 and α3, the MIM peptides have opposite orientations (N→C). (F) Superposition of canonical MIM2 of CHMP6 (orange C^α^ trace, PDB ID 2K3W) (Kieffer et al., 2008) and MIM1 motif of CHMP1A (aqua C^α^ trace, PDB ID 2JQ9) (Stuchell-Brereton et al., 2007) onto the predicted structure of pUL71 (violet C^α^ trace) in complex with human VPS4 (cyan ribbons). MIM2 groove residues V13 (*left*) and MIM1 groove residue L64 (*right*) highlighted as red sticks. (G) Co- transfection of pUL71 with FLAG-tagged VPS4A, either full-length (WT), lacking the N-terminal 84 residues encoding the MIT domain (ΔMIT), or with single amino acid substitutions that prevent binding to MIM2 (V13D) or MIM1 (L64D) regions. Nuclei are outlined in single channel images and shown (DAPI, blue) in merge. Scale bars = 10 µm.

To further probe the predictive power of the AlphaFold-Multimer model, umbrella sampling (Lemkul and Bevan, 2010) was used to computationally probe how structurally-informed mutations effect the binding of pUL71 to the VPS4A MIT domain. Briefly, the model of VPS4A bound to wild-type or mutant pUL71(300–325) was taken as the starting structure and a series of conformations were generated along a trajectory of increasing centre-of-mass (COM) between pUL71(300–325) and VPS4A MIT using steered molecular dynamics. Each of these conformations was then simulated for 10 ns, retaining the COM distance using a biasing function, to give an ensemble of structures sampling the pUL71(300– 325):VPS4A complex as a function of increasing COM distance (Fig. 4 Supp. 1). A curve of potential of mean force (PMF) as a function of the COM distance was generated using the maximum likelihood weighted histogram analysis method (Doudou et al., 2009; Kumar et al., 1992), allowing estimation of the free energy of binding (ΔG_bind_) as the difference between the PMF of the bound and unbound state (Lemkul and Bevan, 2010). pUL71 mutations known to inhibit VPS4A binding (P315A+P318A[PPAA] and V317D, Fig. 1D,E,G) were tested, as were mutations predicted to be deleterious via removal of a hydrophobic side chain (P315A, P318A) or replacement of a hydrophobic interacting residue with a bulky charged residue (I307R, M311R). The mutation R312E was also tested as a control, as inverting the side chain charge of this residue that faces away from VPS4A in the predicted complex structure should not affect pUL71 binding (Fig. 3A). Results of the umbrella sampling are shown (Fig. 4A,B). As anticipated, the calculated ΔG_bind_ of the pUL71(R312E) peptide did not differ significantly from the wild-type peptide (ΔΔG_bind_ ≈ 0; Fig. 4C). Similarly, peptides with the pUL71(V317D) and pUL71(PPAA) control mutations had less negative (weaker) calculated free energies of binding (ΔΔG_bind_ > 0; Fig. 4C). The structurally-informed pUL71(I307R) and pUL71(M311R) peptides were similarly predicted to have reduced binding to VPS4A MIT (ΔΔG_bind_ > 0; Fig. 4C), and ITC confirmed that the pUL71(300–325) I307R peptide lacks the ability to bind the VPS4A MIT domain *in vitro* (Fig. 4D). However, the Umbrella analysis suggested that the single hydrophobic side chain mutations pUL71(P315A) and pUL71(P318A) retained the ability to efficiently bind the VPS4A MIT domain (ΔΔG_bind_ > 0; Fig. 4C), despite binding being lost when both mutations are combined in pUL71(PPAA). Immunoprecipitation of co-transfected pUL71 and VPS4A-FLAG confirmed this surprising result (Fig. 4E). The ability of molecular dynamics analysis to accurately predict VPS4A-binding behaviours of individual amino acid substitutions in the pUL71 peptide confirms the high quality of the pUL71(300–325):VPS4A MIT structural model. We thus conclude that pUL71 possesses a viral MIM2-like motif (vMIM2) that adopts a helix plus extended confirmation and binds between helices α1 and α3 of the VPS4A MIT domain.

**Figure 4.**
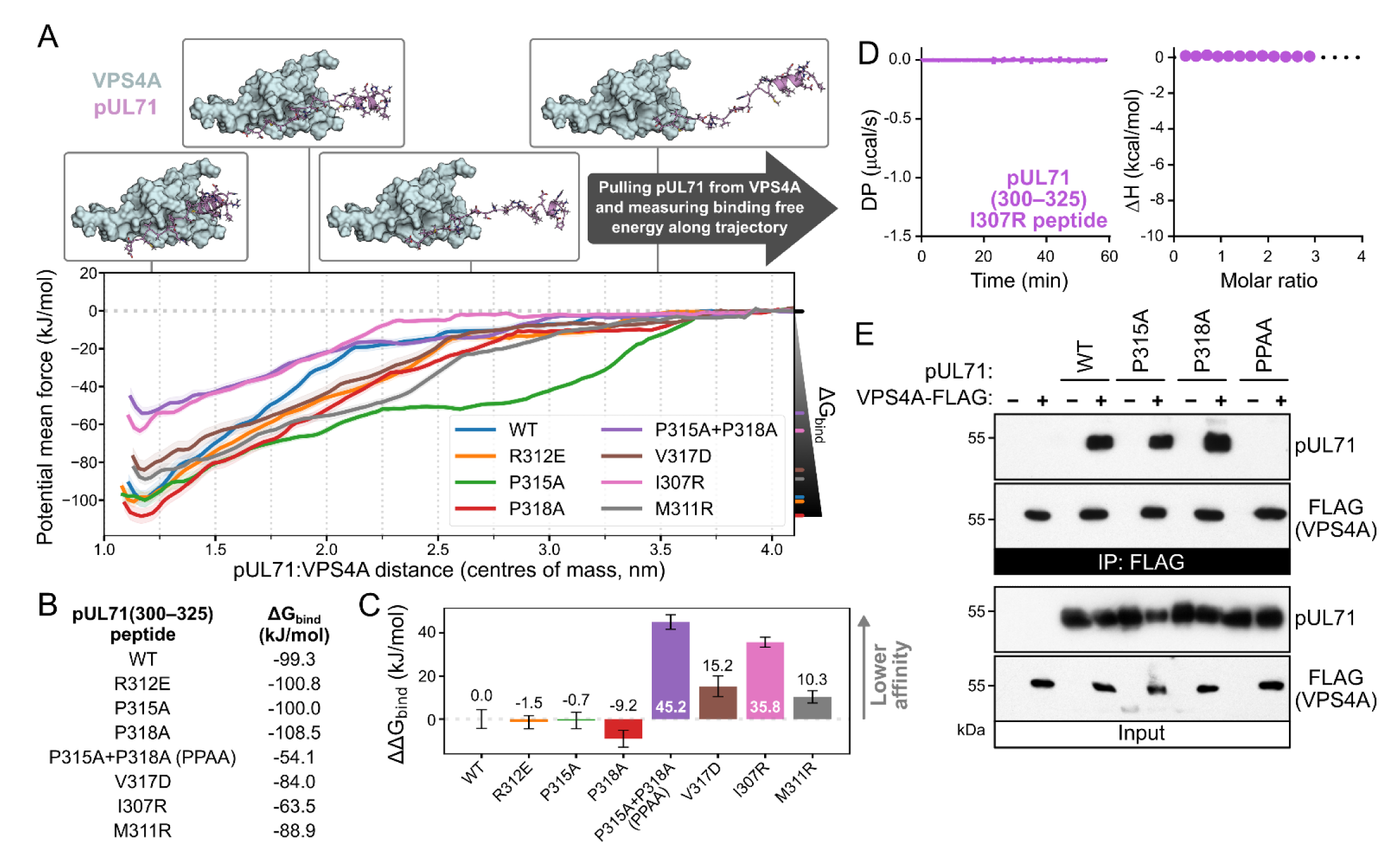
The pUL71(300–325):VPS4A MIT domain structural model has predictive power. (A) Umbrella sampling molecular dynamics (MD) simulations of WT and mutant pUL71(300–325) in complex with the VPS4A MIT domain. For each, potential of mean force ± SD (shaded) from 200 bootstraps of the analysis is plotted as a function of centre of mass between the two polypeptides. The calculated Gibbs free energy of binding (ΔG_bind_) is the difference between the potential mean force minimum (bound) and maximum (unbound) values. (B) ΔG_bind_ for pUL71(300–325) WT and mutants. (C) Difference in ΔG_bind_ for pUL71(300–325) mutants compared to WT (ΔΔG_bind_; mean ± SD for 200 bootstraps). Positive ΔΔG_bind_ values indicate reduced binding affinity, negative values indicate increased affinity. (D) ITC analysis demonstrating a lack of binding when a peptide corresponding to pUL71 residues 300–325 with an I307R substitution is titrated against the VPS4A MIT domain. (E) Anti-FLAG IP from cells co-transfected with VPS4A-FLAG and WT, P315A, P318A, or P315A+P318A (PPAA) pUL71. Samples were immunoblotted using antibodies as shown.

### VPS4A binding is conserved amongst cytomegaloviruses and human β-herpesviruses

While multiple alignment of β-herpesvirus homologue sequences failed to identify conservation of pUL71 residues 300–325 (Fig. 1A), manual inspection of the C-terminal regions of various primate cytomegaloviruses indicated that each might contain a VPS4A-binding vMIM2 (Fig. S3A). In particular, the fourth (P_4_) residue of the ϕ-LP-VPS-xLP MIM2 consensus (Kieffer et al., 2008) is absolutely conserved, as is P_7_, and there is conservation of a hydrophobic residue (A, L, P or V) at the sixth and tenth positions. AlphaFold-Multimer structure prediction of the MIM2-like regions indicates that each is likely to adopt a helix-plus-extended confirmation to bind the MIT domains of VPS4A from their cognate host species between helices α1 and α3, as seen for HCMV pUL71 (Fig. S3B–D). AlphaFold-Multimer analysis of mouse cytomegalovirus (MuHV-1) and the England isolate of rat cytomegalovirus (MuHV-8) suggests that these viruses will also bind the VPS4A MIT domains of their cognate hosts (Fig. S4). Immunoprecipitation experiments confirm that mouse cytomegalovirus binds FLAG-tagged human VPS4A (Fig. S4), the MIT domains of mice and men sharing 97.6% sequence identity.

Beyond non-human cytomegaloviruses, careful inspection of the C-terminal tails identified potential vMIM2s in pUL44 of human β-herpesvirus HHV6A and HHV6B (which have identical sequences in this region) and HHV7 (Fig. 5A). AlphaFold-Multimer confidently predicts HHV6 pU44(174–199) and HHV7 pU44(164–189) to bind the VPS4 MIT domain in a very similar conformation to HCMV pUL71 (Fig. 5B), although the orientation of the N-terminal helix is less confidently predicted (lower pLDDT, higher PAE) and differs slightly between HCMV and HHV6/7. Sequence alignment of the potential vMIM2s shows that pUL71 I307, which binds a hydrophobic surface on VPS4A (Fig. 3A) and is important for binding (Fig. 4A–D), is replaced by glutamate in HHV6 and HHV7 pU44. However, close inspection of the structural models suggests structural rearrangement of the side chains that bury this hydrophobic surface: the side chains of Y186 and L189 of HHV6 pU44 (Y176 and L179 in HHV7) occupy similar space to pUL71 side chains I307 and M311 (Fig. 5B), with L189 having a similar orientation as observed for L170 in the structure of the CHMP6 canonical MIM2 bound to VPS4A (Kieffer et al., 2008). While the position of hydrophobic residues within the helical region that binds VPS4A is flexible, pUL71 homologues conform to a vMIM2 consensus sequence <u>Ωxx</u>xPxϕϕxxxϕ, where Ω denotes a large hydrophobic residue, x denotes any residue, ϕ denotes a small hydrophobic residue (including proline), and where the underlined residues are within an α-helix.

**Figure 5.**
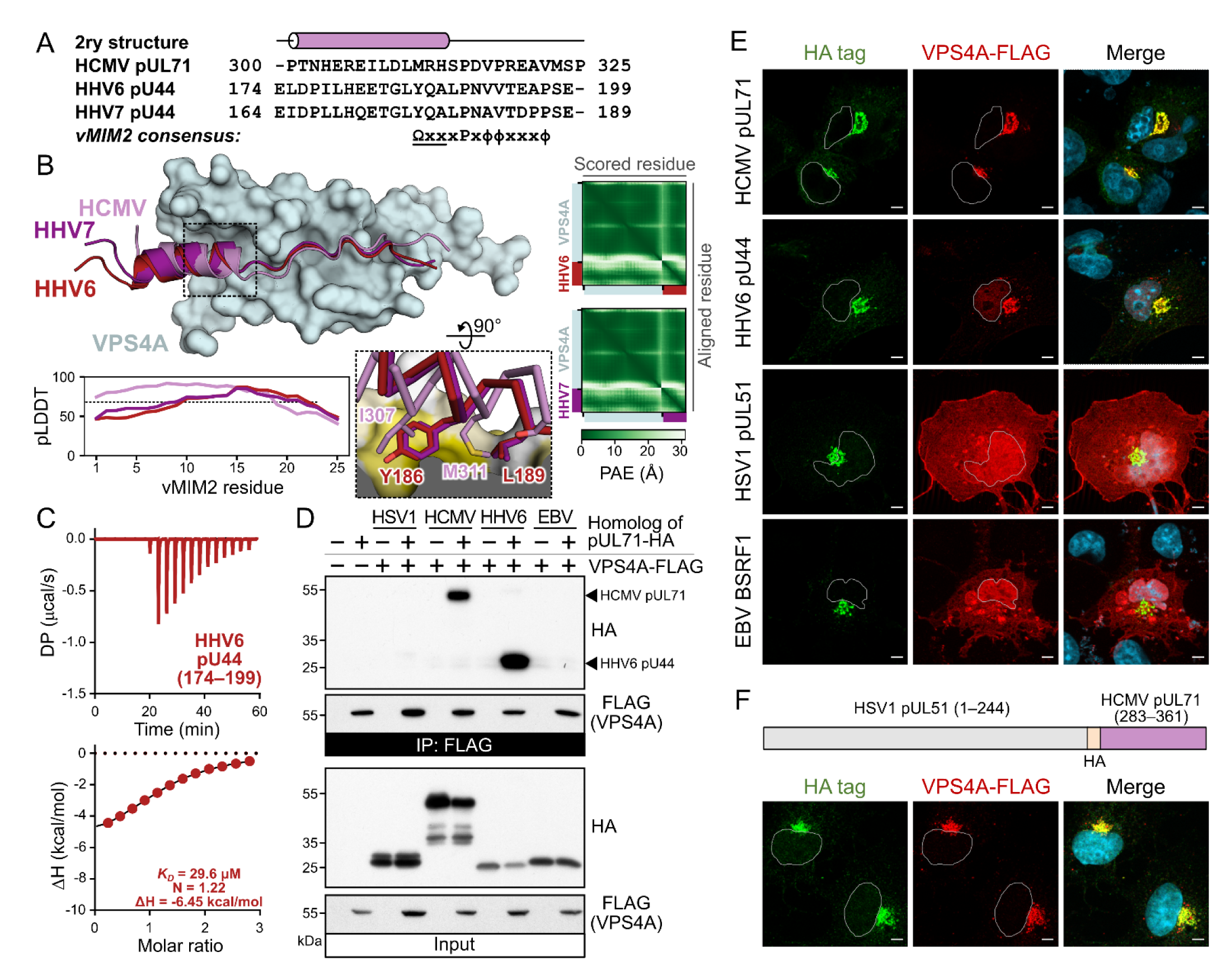
VPS4A binding is conserved across human β-herpesviruses, but not α- or γ-herpesviruses. (A) Alignment of vMIM2 regions of pUL71 and its homologue (pU44) in HHV6 and HHV7. Note that the displayed region of pU44 has an identical sequence in HHV6A and HHV6B. The predicted secondary structure of pUL71 is shown above. The pUL71 homologue vMIM2 consensus sequence is shown below, where Ω denotes a large hydrophobic residue, x denotes any residue, ϕ denotes a small hydrophobic residue (including proline), and where the underlined residues are within an α-helix. (B) Predicted structure of residues from the vMIM2 of β-herpesvirus pUL71 homologues (violet, red and purple ribbon for HCMV, HHV6 and HHV7, respectively) in complex with the VPS4A MIT domain (cyan molecular surface). *Inset:* Predicted interactions between hydrophobic residues of the pUL71 homologue helical regions and the VPS4A MIT domain. VPS4A molecular surface is coloured by residue hydrophobicity from white (polar) to yellow (hydrophobic). C^α^ trace and selected side chains of vMIM2s are shown. *Below:* Per-residue pLDDT scores of the vMIM2s, with values above 70 (dashed line) representing regions predicted with high confidence. *Right:* PAE matrices for predicted HHV6 and HHV7 complexes. (C) ITC analysis of the VPS4A MIT domain binding a peptide corresponding to the HHV6 pU44 vMIM2 (residues 174–199). (D) Anti-FLAG IP from cells co-transfected with VPS4A-FLAG and HCMV pUL71 or homologues from HSV-1 (pUL51), HHV6 (pU44) and EBV (BSRF1). Samples were immunoblotted using antibodies as shown. (E) Co-transfection of VPS4A-FLAG and HCMV pUL71 or homologues from HSV-1 (pUL51), HHV6 (pU44) and EBV (BSRF1). (F) Co-transfection of VPS4A-FLAG with a chimeric construct encoding full length HSV-1 pUL51 followed by an HA tag plus the C-terminal region of HCMV pUL71 (residues 283–361, which includes the vMIM2). Nuclei are outlined in single channel images and shown (DAPI, blue) in merge. Scale bars = 10 µm.

ITC analysis shows that HHV6 residues 174–199 bind purified human VPS4A MIT domain with 30.9 ± 5.76 µM affinity (±SD, n=3; Fig. 5C and Table 1). The C-terminal tails of HSV-1 pUL51 and EBV BSRF1, α- and γ-herpesvirus homologues of pUL71, respectively, are also predicted to be disordered and both proteins associate with juxtanuclear membranes (Butt et al., 2020; Nozawa et al., 2003; Yanagi et al., 2019). Careful inspection of the HSV-1 pUL51 and EBV BSRF1 C-terminal sequences failed to reveal a vMIM2 sequence. Consistent with this, immunoprecipitation of transfected C-terminally HA-tagged HCMV pUL71 or HHV6 pUL44 showed that these pUL71 homologues bind efficiently to co-transfected VPS4A-FLAG in cultured cells, while HSV-1 pUL51 and EBV BSRF1 do not (Fig. 5D). Similarly, VPS4A- FLAG is not recruited to juxtanuclear membranes when co-transfected with pUL51 or BSRF1, while it is recruited by both pUL71 and HHV6 pUL44 (Fig. 5E). To confirm that the lack of association in these homologues arose from an absence of interaction with VPS4A, an HA epitope tag and the C-terminal tail of pUL71, including the vMIM2 (residues 283–361), was appended to the C terminus of pUL51. This chimeric pUL51-HA-pUL71(283–361) protein gained the ability to recruit co-transfected VPS4A- FLAG to juxtanuclear membranes (Fig. 5F). These results confirm that the vMIM2 is conserved across human β-herpesviruses, but not in α- or γ-herpesviruses, and that this motif is sufficient to confer recruitment of VPS4A to biological membranes.

### Accumulation of VPS4A at the cytoplasmic viral assembly compartment requires pUL71

Localisation of VPS4A to the cytoplasmic viral assembly compartment (cVAC) during HCMV infection has been shown previously (Das and Pellett, 2011; Dell’Oste et al., 2014; Tandon et al., 2009). We sought to recapitulate these findings and investigated whether accumulation of VPS4A at the cVAC is a general feature of HCMV infection. Human fibroblasts transiently expressing VPS4A-FLAG from a Tet-inducible promoter were infected with different HCMV strains and VPS4A-FLAG expression was induced at 1 day post infection (dpi) by addition of doxycycline. Detection of HCMV tegument protein pUL71 served as a control for infection and furthermore served as marker for the perinuclear cVAC, as localisation of pUL71 to the cVAC was previously shown (Dietz et al., 2018; Read et al., 2019; Womack and Shenk, 2010). We detect accumulation of VPS4A-FLAG at the area of the cVAC in all infected cells and for the various HCMV strains, including clinical isolates (Fig. 6A). Furthermore, there is noticeable overlap of VPS4A-FLAG signals with the signals for pUL71, which is consistent with our results from transient expression experiments, indicating an ability of HCMV pUL71 to interact with VPS4A in the context of infection that results in recruitment of VPS4A to the cVAC and thus the site of secondary envelopment. To further corroborate these results, bimolecular fluorescence complementation (BiFC) was performed in which the N-terminal half of Citrine (YN) was fused to the C terminus of pUL71 and expressed from the endogenous locus of a recombinant strain of HCMV TB40/E (TB71-YN), while the C-terminal half of Citrine (YC) was provided by transfection with vectors encoding YC-VPS4A-FLAG and mutants in the VPS4A MIT domain under the control of doxycycline (Fig. 6D). Citrine fluorescence was detected in infected cells that also expressed YC-VPS4A-FLAG and YC- VPS4A-FLAG-L64D, but not for YC-VPS4A-FLAG-V13D. These data are consistent with our previous results from transient expression and show that in infected cells VPS4A accumulates at the cVAC via an interaction with pUL71 that depends on an interaction of the VPS4A MIT domain with the vMIM2.

**Figure 6.**
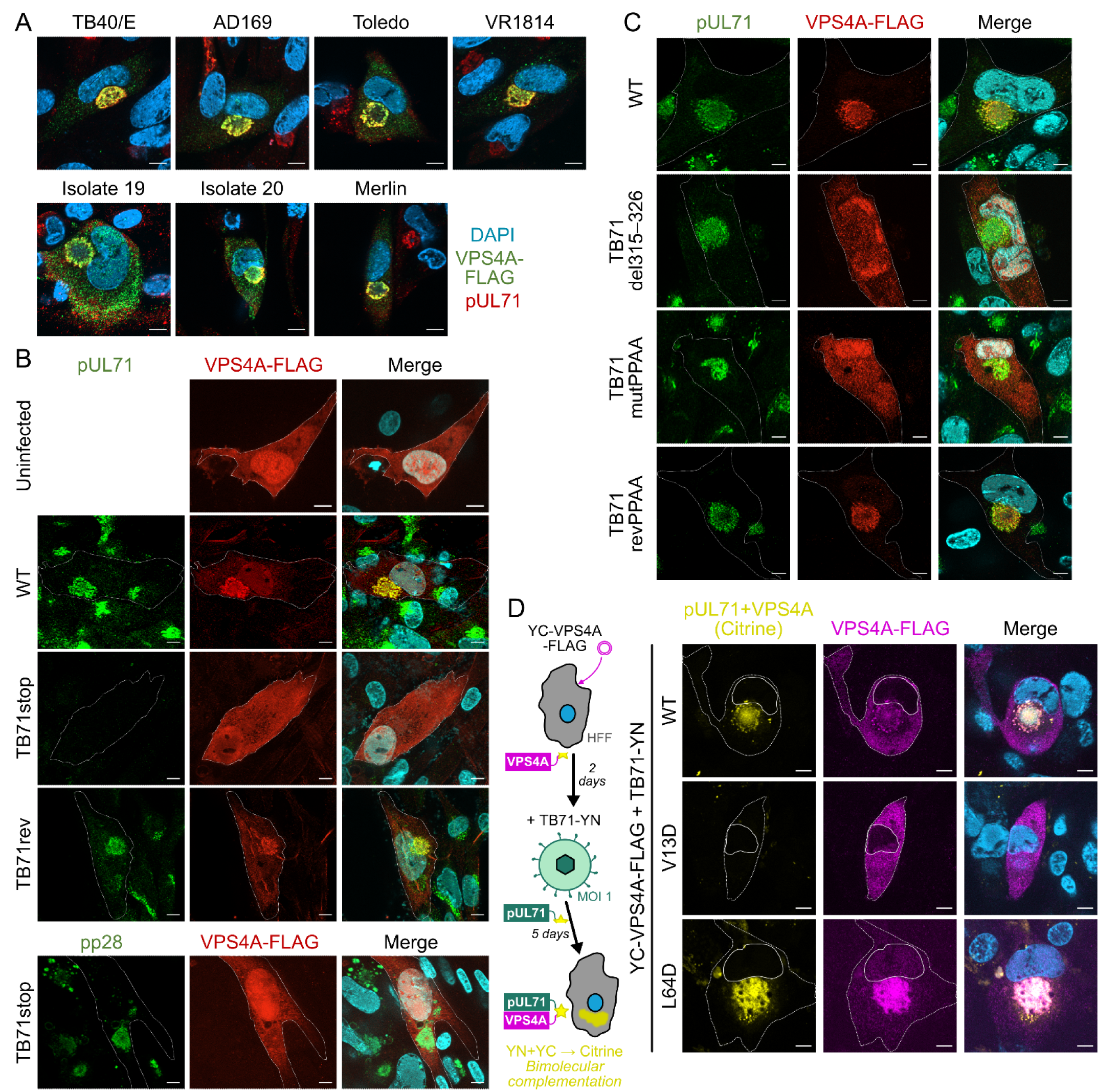
pUL71 vMIM2 is required for VPS4A recruitment to the cVAC during infection. (A) MRC-5 cells transiently expressing VPS4A-FLAG from an inducible expression vector were infected (MOI 0.5) with the indicated strains of HCMV. Expression of VPS4A-FLAG was induced 1 dpi by addition of doxycycline and intracellular distribution VPS4A-FLAG was examined at 5 dpi via antibody detection of the FLAG epitope. The cVAC is denoted by additional staining for HCMV pUL71 (red) and nuclei are shown (DAPI, blue). (B, C) MRC-5 cells transiently expressing VPS4-FLAG were mock-infected or infected (MOI 0.5–1) with indicated viruses. Expression of VPS4A was induced 1 dpi and cells were fixed and immunostained with the antibodies shown at 5 dpi. Selected cells are outlined in single channel images and nuclei are shown (DAPI, cyan) in merge. (B) Cell were infected with WT HCMV, with a recombinant virus lacking pUL71 expression (TB71stop), or the revertant virus with restored pUL71 expression (TB71rev). (C) Cells were infected with WT HCMV, with recombinant virus lacking the vMIM2 motif (TB71del315–326), with mutations P315A+P318A in the vMIM2 to disrupt VPS4A binding (TB71mutPPAA), or with the wild-type sequence subsequently restored (TB71revPPAA). (D) Bimolecular fluorescence complementation using split Citrine (residues 1–173, YN, and residues 156– 239, YC) confirms a physical interaction between pUL71 and VPS4A during infection. MRC-5 cells conditionally expressing WT, MIM2-binding groove (V13D) or MIM1-binding surface (L64D) mutant human VPS4A that was N- and C-terminally tagged with YC and FLAG, respectively, were infected (MOI 1) with HCMV strain TB40/E where pUL71 was C-terminally tagged with YN (TB71-YN). Expression was induced 2 dpi by addition of doxycycline. Cells were fixed, immunostained (FLAG, magenta), and Citrine fluorescence from reconstitution of its constituent parts (yellow) was visualised at 5 dpi. Selected cells and their nuclei are outlined in single channel images are nuclei are shown (DAPI, blue) in merge. Scale bars = 10 µm.

Next, we investigated the role for pUL71 in VPS4 accumulation during infection by testing previously generated and characterised recombinant HCMV, TBstop71 (unable to express pUL71) and TBrev71 (expression of pUL71 repaired) (Schauflinger et al., 2011). Recruitment of VPS4A-FLAG to the perinuclear cVAC failed in cells infected with the TBstop71 virus, but was detectable in infection with the parental virus and TBrev71 revertant (Fig. 6B). In summary, these data show that the recruitment of VPS4A to the cytoplasmic site of virion assembly in the course of infection depends on pUL71.

### Recruitment of VPS4A during infection by pUL71 requires the vMIM2

To investigate whether the interaction of pUL71 with VPS4A is involved in virion morphogenesis or modulates the function of pUL71 during secondary envelopment, recombinant viruses carrying mutations of the vMIM2 were generated by markerless BAC mutagenesis. We generated two recombinant viruses in which the interaction of pUL71 with VPS4A is disrupted: (i) a vMIM2 deletion mutant, TB71del315–326, and (ii) a double point mutant of the two proline residues (P315, P318) of the vMIM2 to alanine residues (Fig. 1D,E,G), resulting in mutant TB71mutPPAA. A revertant (YB71revPPAA) was generated by mutating TB71mutPPAA to restore the wild-type sequence.

These viruses were used in infection experiments to observe the recruitment of VPS4A-FLAG in immunofluorescence analysis. While VPS4A was recruited to the cVAC in cells infected with wild-type and TB71revPPAA viruses, VPS4A remained distributed throughout the entire cell and failed to be recruited to the site of pUL71 localisation in cells infected with the interaction-deficient mutant viruses TB71del315–326 and TB71mutPPAA (Fig. 6C). This shows the successful generation of the intended recombinant viruses and that the vMIM2 in the C terminus of pUL71 is necessary to recruit VPS4A to the cVAC in infection.

To investigate the importance of the mutation of the VPS4A interaction motif of pUL71 in HCMV infection, the replication kinetics of recombinant viruses TB71mutPPAA, TB71revPPAA (revertant virus of TB71mutPPAA) and TB71del315–326 in HFFs were investigated in single step kinetics using an MOI of 3 and in multi-step kinetics using an MOI of 0.01. The virus yields in the supernatants of infected cells, harvested at the indicated times, were determined by titration on HFFs. The growth curves of virus mutants with mutated VPS4 interaction motif (TB71mutPPAA and TB71del315–326) were comparable to those of HCMV wild-type virus (Fig. 7A). Cell-associated viral spread, assessed for TB71mutPPAA, was also identical to that of wild-type virus (Fig. 7B), suggesting overall that the generation and release of infectivity, and cell-to-cell spread, is not affected by mutation of the VPS4A interaction motif of pUL71.

**Figure 7.**
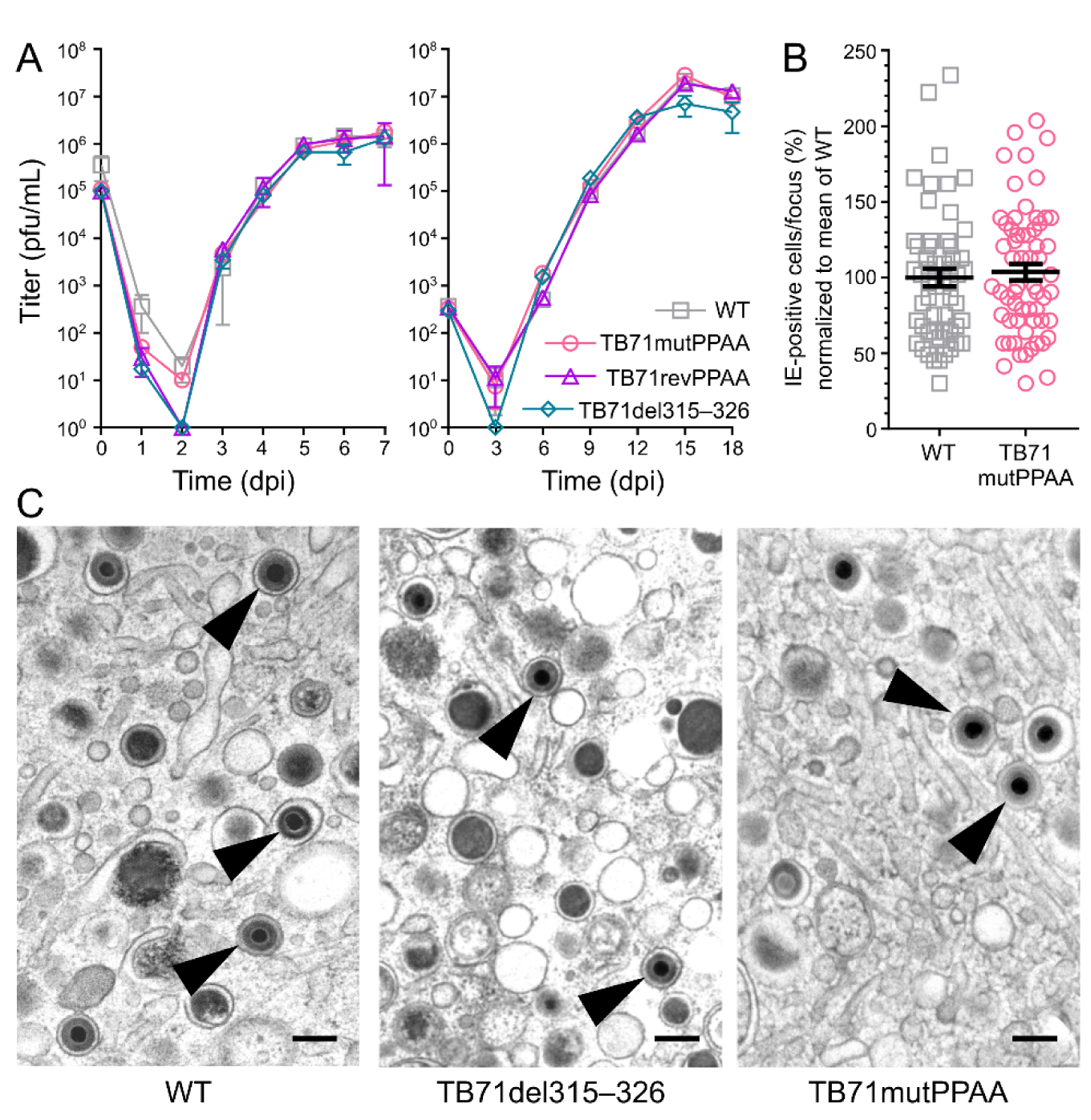
VPS4A recruitment to the cVAC is not required for efficient virus wrapping or spread in human fibroblasts. (A) Mutations in the pUL71 vMIM2 motif do not affect virus release. For single- step kinetics (*left*), HFFs were infected (MOI 3) with HCMV WT (□), TB71mutPPAA (○) and TB71revPPAA (Δ) or TB71del315–326 (◊). For multi-step kinetics (*right*) cells were infected at MOI 0.01. The supernatant of infected cells was harvested at the indicated times post infection and the virus yield was determined by titration on HFFs. Mean ± SD is shown (n = 4; two independent experiments performed in technical duplicate). Virus yields of the inocula are given at time zero. (B) Relative size of plaques formed by HCMV WT and TB71mutPPAA after 9 days of infection in a focus expansion assay. Infected cells were detected by immunostaining for HCMV IE1/2 protein. Each data point represents the relative number of IE-positive nuclei per focus. Mean ± SEM (n = 56–61 foci) is shown for each virus (black line) normalised to the mean focus size for WT. (C) Electron micrographs showing representative areas of the cVAC at 5 dpi from cells infected with WT HCMV or with the indicated mutant viruses. Arrow heads mark fully enveloped virus particles. Scale bar = 200 nm.

### Mutation of the pUL71 vMIM2 does not affect the ultrastructural phenotype of HCMV infected cells

The influence of mutations of the vMIM2 in pUL71 on the function of pUL71 for HCMV secondary envelopment was investigated by transmission electron microscopy (TEM) of infected cells at 5 dpi. Previous studies have shown an involvement of pUL71 in secondary envelopment (Schauflinger et al., 2011; Womack and Shenk, 2010). In addition to an increased number of nucleocapsids at different budding stages, cells infected with HCMV lacking pUL71 exhibit enlarged multivesicular bodies, many of which are in close proximity to the cVAC, and budding processes can be observed at these enlarged vesicular structures (Schauflinger et al., 2011).

The ultrastructural phenotype of both interaction-deficient mutant viruses, TB71mutPPAA and TB71del315–236, was undistinguishable to that of wild-type virus infected cells (Fig. 7C). An enlargement of MVBs or other vesicles as previously described in absence of pUL71 were not found. Furthermore, there was no defect in secondary envelopment apparent. Quantification of cytoplasmic morphogenesis stages of capsids at the cVAC confirmed that the majority of virus particles were enveloped to comparable degrees in the wild type and recombinant viruses (Table 2). In summary, these data show that recruitment of VPS4A to the cVAC by pUL71 is dispensable for viral morphogenesis and growth in cultured cells.

**Table 2.** Quantification of HCMV secondary envelopment. The cVAC of HFF cells 5 days post-infection with wild type, TB-71mutPPAA, and TB-71del314-326 HCMV was analysed by electron microscopy (Figure 7). Relative numbers represent percent of enveloped particles, non-enveloped particles attached to membranes (budding particles), and non-enveloped particles (naked particles) observed.

| Virus | No. of cells analysed | No. of particles analysed | mean % particles ± SD | | |
| --- | --- | --- | --- | --- | --- |
|  |  |  | Enveloped particles | Budding particles | Naked particles |
| Wild type | 24 | 888 | **87.58** ± 11.26 | 11.97 ± 11.01 | 0.46 ± 1.76 |
| TB-71del315–326 | 14 | 494 | **82.51** ± 10.76 | 17.11 ± 9.89 | 0.37 ± 1.41 |
| TBmutPPAA | 28 | 749 | **74.78** ± 13.08 | 21.88 ± 10.23 | 3.34 ± 5.79 |

## Discussion

While many viral proteins have been identified as important for HCMV secondary envelopment, the molecular interactions between these proteins and with cellular partners that drive wrapping of tegument-covered capsids and the final step of membrane scission remain unclear. Homologues of HCMV pUL71 are conserved across herpesviruses (Lenk et al., 1997; Mocarski Jr., 2007), stimulate virus secondary envelopment (Albecka et al., 2017; Jiang et al., 2017; Klupp et al., 2005; Nozawa et al., 2005; Schauflinger et al., 2011), their N-terminal helical domains have structural homology to cellular ESCRT-III components like CHMP4B (Butt et al., 2020; He et al., 2020), and they are predicted to undergo N-terminal palmitoylation (Nozawa et al., 2003; Roller et al., 2014) that is reminiscent of the N-terminal myristoylation of CHMP6 (Yorikawa et al., 2005). We show here that the homology of pUL71 to ESCRT-III components extends further in the β-herpesviruses: The C-terminal tail of HCMV pUL71 has a vMIM2 that binds directly to the MIT domain of VPS4A (Fig. 1,2). This motif is conserved across pUL71 from other human β-herpesviruses (Fig. 5) and across cytomegaloviruses of other species (Fig. S3, S4). We are unaware of any prior reports of a virus encoding a MIM or directly binding the VPS4A MIT domain.

The pUL71 vMIM2 is necessary and sufficient to recruit VPS4A to specific membranes in co-transfected cells (Fig. 5) and to sites of virus assembly in HCMV infection (Fig. 6), but it is dispensable for virus replication and secondary envelopment (Fig. 7). The lack of a defect in secondary envelopment or viral growth in viruses lacking a functional pUL71 vMIM2 (TB71del315–326 and TB71mutPPAA) is consistent with a recent report that VPS4 activity is not necessary for HCMV replication (Streck et al., 2018). This study showed that single-step replication of HCMV is not impaired by expression of dominant negative mutants of VPS4A (E228Q) or ESCRT-III components CHMP4B and CHMP6, and the morphology of mature virions and wrapping compartments are unchanged in the presence of these proteins. There is a mild defect in virus spread in the presence of dominant negative CHMP4B and CHMP6, quantitated via immunocytometry of cells expressing immediate early proteins following low MOI infection, but the effect of dominant negative VPS4 on spread was not analysed. We observe no difference in infectious particle production during multi-step growth curves and focus expansion assays of WT versus vMIM2 mutants of HCMV (Fig. 7A), suggesting that if there is any spread difference in HFFs it is likely to be very minor.

If VPS4 activity is not required for secondary envelopment, then why is the ability to bind the VPS4A MIT domain conserved across β-herpesviruses? The VPS4A MIT domain binds the isolated pUL71 vMIM2 more tightly than it does the MIM2 of CHMP6 (*K*_D_ = 2.3 µM versus 11.2 µM; Fig. 2 and Table 1). The HHV6 pU44 vMIM2 binds less tightly (*K*_D_ = 30.9 µM), but high abundance of pUL71 and homologues at later stages of infection (Weekes et al., 2014) and their potential ability to polymerise (Butt et al., 2020) would allow them to compete effectively with endogenous CHMP6 for VPS4A binding. It is thus possible that pUL71-mediated recruitment to the cVAC *sequesters* VPS4A, preventing it from having anti-viral effects elsewhere in the cell, or *redirects* it to support pro-viral functions. Fraile-Ramos and colleagues (Fraile-Ramos et al., 2007) reported that simultaneous siRNA depletion of VPS4A and VPS4B from RPE1 cells increased HCMV strain RCMV288 particle production, as did treatment of the cells with proteases leupeptin plus E64, and they hypothesised that the cellular ESCRT-III machinery may promote HCMV budding into MVBs that are delivered to the lysosome for destruction. We do not observe an increase in virus growth in either single-step or multi-step growth curves, suggesting that inhibition of VPS4 activity is not beneficial for virus growth in cultured human fibroblasts. However, it remains possible that such inhibition of virion degradation has a more potent effect upon virus growth in other cells that HCMV infects such as monocytes or in terminally differentiated cells that have exited the cell cycle. Additionally, MHC-II molecules are known to acquire their peptide cargos when present on intraluminal vesicles within the endosomal system of ‘professional’ antigen presenting cells such as macrophages (Pishesha et al., 2022). It may be that inhibition of ESCRT-III activity, which has been postulated to promote budding of MHC-II into these intraluminal vesicles (Neefjes et al., 2011), could represent a mechanism by which HCMV evades MHC- II mediated host immunity. Alternatively, pUL71 might redirect VPS4A activity to stimulate budding of non-virus vesicles into the lumen of endolysosomal compartments. HCMV infection is known to stimulate the production of extracellular vesicles (EVs) that enhance virus spread (Streck et al., 2020) and EVs from virus-infected cells are enriched for both pUL71 and its direct binding partner pUL103 (Butt et al., 2020; Turner et al., 2020). pUL71-driven VPS4A recruitment may thus stimulate secretion of a specific repertoire of EVs from infected cells, budding either at the cVAC or at other sites within the infected cell, to promote spread or otherwise modulate the infected-cell environment.

A second hypothesis is that the vMIM2 of pUL71 alters the trafficking and secretion of virus particles. While it is well established that individual HCMV virions can bud into vesicles that are transported to specific regions of the cell surface for release (Schauflinger et al., 2013, 2011), recent studies have shown that HCMV particles can also accumulate in MVB-like compartments before being released into to the extracellular milieu (Flomm et al., 2022) and that components of the cellular exosome biogenesis machinery may contribute to virion maturation (Turner et al., 2020). ESCRT machinery and VPS4 activity are well established as key components of MVB morphogenesis and it is possible that pUL71 influences the pathway of HCMV particle release by altering the generation/maturation of MVBs via redirecting VPS4A activity. A role for pUL71 in defining the pathway of exit for virus particles is supported by the observation that TB40-71stop infection causes the accumulation of enlarged virus-containing MVBs in the proximity of the cVAC (Schauflinger et al., 2011). We did not observe similar alterations to the MVB morphology of cells infected with HCMV expressing pUL71 without a vMIM2 (TB71del315–326 and TB71mutPPAA), suggesting that redirection/sequestration of VPS4A is not sufficient for this phenotype and that other activities of pUL71 must also be involved.

There are over 20 human proteins with MIT domains, including five ATPases, and seven different modes of interaction between short linear motifs and MIT domains have been structurally characterised (Wenzel et al., 2022). Interactions with MIM2 are generally weaker than interactions with other MIMs (e.g. MIM1 and MIM3) and only a very limited subset of MIT-domain containing proteins, including just two ATPases (VPS4A and VPS4B), bind isolated MIM2 regions with appreciable affinity (Wenzel et al., 2022). The other MIM2-binding proteins are involved in protein deubiquitylation (AMSH and USP8) (McCullough et al., 2004; Naviglio et al., 1998; Row et al., 2007), the midbody abscission checkpoint (MITD1) (Agromayor et al., 2009), membrane trafficking (SNX15) (Phillips et al., 2001) or have a pseudo-kinase fold (RPS6KC1) (Hayashi et al., 2002). It is possible that one of these proteins could represent the functional target of pUL71, although this awaits experimental investigation. While the functional consequences of VPS4A recruitment by pUL71 remain enigmatic, the presence of a conserved C-terminal vMIM2 adds additional circumstantial evidence to support the hypothesis (Streck et al., 2018) that pUL71 may have ESCRT-III–like activity. How HCMV deploys this to remodel intracellular membranes and produce new virus particles remains to be shown.

## Methods

### Sequence analysis and structure prediction

For sequence conservation analysis, homologues of pUL71 from HCMV strain TB40/E (UniProt C8BKL7) were obtained by performing a BLASTP search of a non-redundant (NCBI clustered NR) sequence database (Sayers et al., 2022; Steinegger and Söding, 2017) with manual inspection to remove incomplete sequences. The 19 representative β-herpesvirus pUL71 homologue sequences were aligned using COBALT (Papadopoulos and Agarwala, 2007) using default parameters. Sequence conservation was calculated using Jalview (Livingstone and Barton, 1993).

Sequences of vMIM2s in β-herpesvirus pUL71 homologues were identified by manual inspection and structures of the VPS4A MIT domain (residues 1–84) in complex with pUL71 and homologue vMIM2s were predicted using AlphaFold-Multimer (Evans et al., 2021) version v2.2.1. The structures of vMIM2s from HCMV pUL71 (UniProt C8BKL7), HHV6A pU44 (UniProt A0A140AK71) and HHV7 pU44 (P52474) were predicted in complex with human VPS4A (UniProt Q9UN37). Sequences of non-human cytomegalovirus pUL71 homologues and VPS4A from their cognate host species were as follows (UniProt IDs in parentheses): simian cytomegalovirus (G8XTD2) with *Chlorocebus sabaeus* VPS4A (A0A0D9QVU0), rhesus macaque cytomegalovirus (I3WFB8) with *Macaca mulatta* VPS4A (A0A1D5R8T0), saimiriine betaherpesivirus 4 (G8XSX5) with *Saimiri boliviensis boliviensis* VPS4A (F6ZNA4.1), cynomolgus macaque cytomegalovirus (G8H1A5) with *Macaca fascicularis* VPS4A (A0A2K5TVF2), chimpanzee cytomegalovirus (Q8QS28) with Pan troglodytes VPS4A (K7BC82), Aotine betaherpesvirus 1 (G8XUD6) with *Aotus nancymaee* VPS4A (A0A2K5CPK0), mouse cytomegalovirus (D3XDP9) with *Mus musculus* VPS4A (Q8VEJ9), and England isolate of rat cytomegalovirus (K7XXY4) with *Rattus norvegicus* VPS4A (Q793F9). Structures were superposed and structural images were prepared using an open-source version of PyMOL (Schrödinger). Numerical analyses were plotted using Matplotlib version 3.3.2 (Hunter, 2007).

### Primers

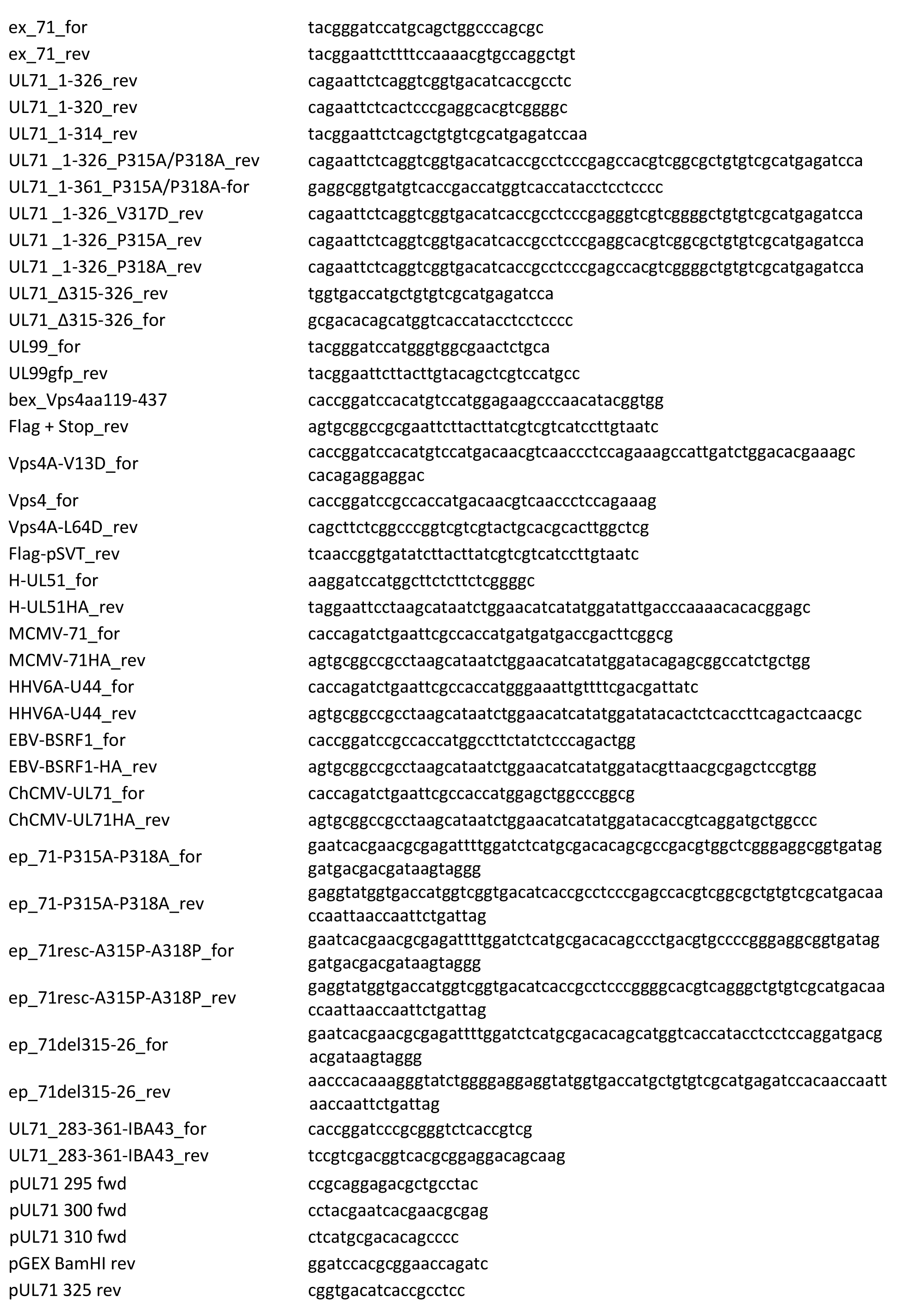
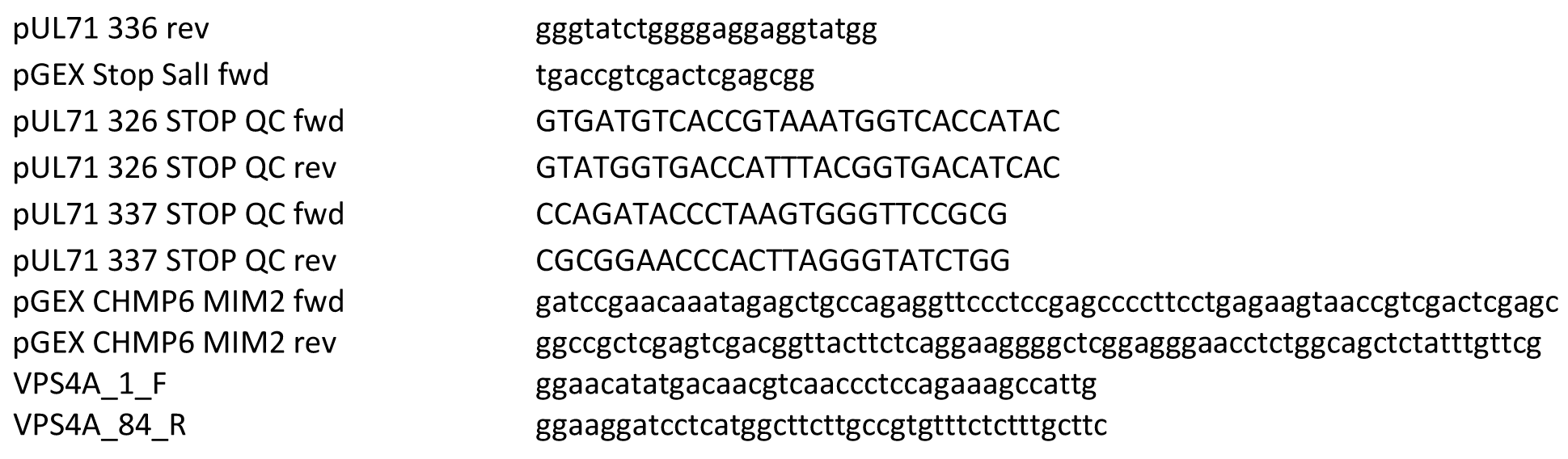

### Viruses

HCMV strains AD169 (Rowe et al., 1956), Toledo (Quinnan et al., 1984), Merlin (Davison et al., 2003), VR1814 (Grazia Revello et al., 2001) and TB40/E (Sinzger et al., 1999) and two clinical isolates (isolates 19 and 20, kindly provided by C. Sinzger, Ulm University Hospital, Germany) were used in this study. This panel was chosen as it covers the most commonly used cell culture passaged strains and two isolates that grow highly cell associated. In addition, bacmid derived viruses were used. Reconstituted virus from bacterial artificial chromosome (BAC) clone TB40-BAC4 of the endotheliotropic HCMV strain TB40/E (accession number EF999921.1.) (Sinzger et al., 2008) served as wild type virus (WT) in this study. Generation and propagation of recombinant viruses, TBstop71 and TBrev71 have been described elsewhere (Schauflinger et al., 2011). Recombinant viruses TB71del315–326, TB71mutPPAA (mutations P315A and P318A) and the revertant thereof TB71revPPAA were generated by markerless BAC mutagenesis (Tischer et al., 2006) from BAC clone TB40-BAC4 by using primers ep_71del315- 26_for and ep_71del315-26_rev, ep_71-P315A-P318A_for and ep_71-P315A-P318A_rev, or ep_71resc-A315P-A318P_for and ep_71resc-A315P-A318P_rev, respectively. The recombinant virus TB71-YN used for bimolecular fluorescence complementation (BiFC) assay, was generated by markerless BAC mutagenesis from BAC clone TB40-BAC4 by inserting the sequences corresponding to a (GGGGS)2-linker sequence, the c-myc-tag and the N-terminal fragment of the yellow fluorescent protein (YFP) variant Citrine, comprising amino acids 1–173 (YN), before the stop codon of the UL71 open reading frame. The YN sequence was first PCR amplified from an YFP-transfer construct (pEP- EYFP-Citrin-in) retrieved from reference (Tischer et al., 2006) by using primers ep_71Cterm-GS- YN173_for and ep_71Cterm-GS-YN173_rev. All recombinant viruses were reconstituted from bacmid DNAs in MRC5 cells, as described in (Read et al., 2019) after sequence verification.

### Plasmids

All eukaryotic expression plasmids for pUL71 and pUL71 homologues were generated in the backbone of pEF1/Myc-His C expression vector (Invitrogen) by cloning using restriction enzymes. The open reading frame (ORF) of full-length UL71 was amplified from the HCMV wild-type BAC TB40-BAC4 using the primers ex_71_for and ex_71_rev and was the basis for all further UL71 mutants. C-terminally truncated variants UL71_1-326, UL71_1-320 and UL71_1-314 were generated using primers ex_71_for and respective reverse primers UL71_1-326_rev, UL71_1-320_rev and UL71_1-314_rev. Point mutants PPAA (P315A+P318A), V317D, P315A and P318A of the vMIM2 of pUL71 were generated by fusion PCR. For the example of the PPAA mutant, a first fragment UL71_1-326-PPAA was amplified with the primers ex_71_for and UL71 _1-326_P315A/P318A_rev. A second fragment UL71_320-361 was amplified by using primers UL71_1-361_P315A/P318A-for and ex_71_rev. The final PCR product UL71-PPAA was amplified by using fragment UL71_1-326-PPAA as template and ex_71_for and fragment UL71_320-361 as primers. UL71-V317D, UL71-P315A and UL71-P318A were generated similarly manner using the primers UL71 _1-326_V317D_rev, UL71 _1-326_P315A_rev and UL71 _1-326_P318A_rev, respectively, to generate the first fragment. Mutant UL71Δ315–326 was generated by using overlapping primers UL71_Δ315-326_rev and UL71_Δ315-326_for together with primers ex_71_for and ex_71_rev, respectively, followed by PCR using both generated PCR fragments and primers ex_71_for and ex_71_rev. For the generation of pp28-EGFP expression vector, the UL99- EGFP sequence was amplified using primers UL99_for and UL99gfp_rev from the bacmid of vTB28g recombinant virus, which was generated and described previously (Chevillotte et al., 2009). Sequences of pUL71 homologues were amplified by PCR from viral DNA, thereby adding a C terminal HA-tag sequence, and restriction cloned into pEF1/Myc-His C expression vector. Primers H-UL51_for and H- UL51HA_rev were used for cloning of HSV-1 pUL51; primers MCMV-71_for and MCMV-71HA_rev for cloning of MCMV M71; primers HHV6A-U44_for and HHV6A-U44_rev for cloning of HHV6 UL71 homologue U44; primers EBV-BSRF1_for and EBV-BSRF1-HA_rev for cloning of EBV UL71 homologue BSRF1.

Human VPS4A-FLAG, C-terminally fused with the FLAG-tag epitope, was expressed from expression vector pBJ5-Vps4Flag (gift from A. Calistri, University of Padova, Italy). Cloning of this vector is described in detail in (Strack et al., 2003). The VPS4-FLAGΔMIT variant, comprising VPS4A residues 119-437, was generated from pBJ5-Vps4Flag using primers bex_VPS4aa119-437 and Flag+Stop_rev. Point mutant VPS4A-FLAG-V13D was amplified using primers Vps4A-V13D_for and Flag+Stop_rev. Variant VPS4-FLAG-L64D was generated by fusion PCR where primers Vps4_for and Vps4A-L64D_rev generated a fragment that was used as forward primer in a second PCR together with primer Flag+Stop_rev to generate full-length VPS4A. All VPS4A variants were cloned using restriction enzymes into backbone vector pEF1/Myc-His C.

For conditional expression of VPS4A-FLAG, the sequence of VPS4A-FLAG was subcloned from pBJ5- Vps4Flag into a single vector expression system (pSVT28, kindly provided by Z. Ruzsics, University Medical Center Freiburg, Germany) by using restriction enzymes. Expression of VPS4A-FLAG in pSVT28 is under the control of doxycycline inducible simian virus 40 (SV40) promoter. Generation of the pSVT28 inducible vector system is described in (Rupp et al., 2005). For the generation of conditional expression vectors for the BiFC assay, the sequence of the C-terminal half of the YFP variant Citrine (YC) comprising amino acids 156–239 and mutation A206K was amplified with primers BiFC-FP-C155R- pSVT_for and BIFC-HA-C155R_rev from vector pUC-SPYCE(MR) (kindly provided by J. Kudla, University of Muenster, Germany) and described in (Walter et al., 2004). The sequence encoding for a (GGGGS)_2_- linker at the 3-prime end was added using PCR. The sequences of VPS4A-FLAG and mutants VPS4A- FLAG-V13D and VPS4A-FLAG-L64D were amplified using primers Vps4_for and Flag-pSVT_rev. The YC fragment and the VPS4A fragments were fused through restriction enzyme digestion followed by ligation and subsequently amplified using primers BiFC-FP-C155R-pSVT_for and Flag-pSVT_rev. The resulting fusion constructs, YC-VPS4A-FLAG, YC-VPS4A-FLAG-V13D, YC-VPS4A-FLAG-L64D, were cloned by restriction enzymes into conditional expression vector pSVT28.

For recombinant protein expression, the C-terminal tail (residues 283–361) of pUL71 from HCMV strain TB40/E was amplified with primers UL71_283-361-IBA43_for and UL71_283-361-IBA43_rev, and then cloned into pGEX4T2 encoding an N-terminal GST tag and thrombin site. Truncated forms were generated by PCR of the entire plasmid except the region to be excluded using KOD HotStart DNA polymerase (Merck) and the forward primers pUL71 295 fwd, pUL71 300 fwd or pUL71 310 fwd with reverse primer pGEX BamHI rev, or forward primer pGEX Stop SalI fwd with reverse primers pUL71 325 rev or pUL71 336 rev, followed by blunt end ligation. Constructs encoding GST-tagged pUL71(300–325) and pUL71(310–336) were generated by QuikChange mutagenesis of plasmids pUL71(300–361) or pUL71(310–361) using the primers pUL71 326 STOP QC fwd and pUL71 326 STOP QC rev or pUL71 337 STOP QC fwd and pUL71 337 STOP QC rev to replace residues 226 or 337 with stop codons, respectively. The MIM2 of human CHMP6 (residues 166–181 of UniProt Q96FZ7) was cloned into pGEX4T2 by ligation of annealed and phosphorylated primers pGEX CHMP6 MIM2 fwd and pGEX CHMP6 MIM2 rev. The MIT domain (residues 1–84) of human VPS4A (UniProt Q9UN37) was cloned into vector pOPT3G (Graham et al., 2013), encoding an N-terminal GST and human rhinovirus 3C cleavage site, using primers VPS4A_1_F and VPS4A_84_R .

### Cell lines and cell culture

Human embryonic kidney cells (HEK293FT, Invitrogen) were maintained in Dulbecco’s modified Eagle medium (DMEM, Gibco/BRL) supplemented with 10% (v/v) foetal calf serum, 2 mM L-glutamine, 100 U of penicillin and 100 µg of streptomycin per mL. African green monkey kidney fibroblast-like cell line (COS-7) was maintained in minimal essential medium (MEM, Gibco/BRL) supplemented with 5% (v/v) foetal calf serum, 2 mM L-glutamine, 100 U of penicillin and 100 µg of streptomycin per mL. Human foreskin fibroblasts (HFFs) were maintained in MEM (Gibco/BRL) supplemented with 10% (v/v) foetal calf serum (Gibco/BRL), 2 mM L-glutamine (Biochrom AG), 100 U of penicillin, 100 µg of streptomycin per mL (both Gibco/BRL), and 1× non-essential amino acids (Biochrom AG). Human embryonic lung fibroblasts (MRC-5, European Collection of Cell Cultures) were maintained in DMEM (Gibco/BRL) supplemented with 10% (v/v) foetal calf serum (Gibco/BRL), 2 mM L-glutamine (Biochrom AG), 100 U of penicillin, and 100 µg of streptomycin per mL (both Gibco/BRL).

### Antibodies

Antibodies that were used in this study to detect viral and cellular proteins were the monoclonal antibody (MAb) recognizing the FLAG-tag epitope anti-Flag (clone M2, Sigma), the MAb against HCMV pp28, anti-pp28 (clone CH19, Santa Cruz Biotechnology (SCBT)) and the MAb 63-37 recognizing HCMV IE1/2 protein (kindly provided by W. Britt, University of Alabama Birmingham, USA). VPS4A was detected with the polyclonal antibody Vps4 (clone H-165, SCBT). Polyclonal antibody anti-HA (A190- 108A, Bethyl laboratories) was used to detect the HA-tag epitope. HCMV pUL71 was detected with a polyclonal anti-pUL71 antibody described elsewhere (Schauflinger et al., 2011). For protein detection secondary anti-mouse or anti-rabbit antibodies conjugated either to Alexa Fluor 488 and 555 (Invitrogen) for immunofluorescence or to horseradish peroxidase (HRP) (Millipore) for immunoblot analysis were used.

### Virus growth analysis

Multistep and single-step growth kinetics experiments were performed as previously described (Read et al., 2019). Briefly, confluent HFFs were infected in duplicate with an MOI of 0.01 for multistep kinetics and an MOI of 3 for single-step kinetics with indicated viruses. To control for the same initial infection, virus yields of the inocula were determined on HFFs by titration. Supernatants were harvested at the indicated times and stored at -80°C until titration. Determination of virus yields in the inocula and the harvested supernatants from the multistep and single-step growth kinetics experiments was performed in triplicate by titration on HFFs as described previously (Schauflinger et al., 2011).

The efficiency of cell-associated spread was determined in a focus expansion assay, as described previously (Dietz et al., 2018). Briefly, confluent HFFs in 12-well plates were infected with 50, 100, and 150 PFU/well of the respective viruses. After 24 h, the cells were washed thoroughly with warm PBS and cultivated under a 0.65% methylcellulose overlay medium until day 9 of infection. Overlay medium was exchanged with fresh overlay medium at day 5 post infection. For detection of infected cells by indirect immunofluorescence staining, the overlay medium was removed, the cells were washed three times with warm PBS and fixed with ice-cold methanol for 10 min at -20°C. HCMV- infected cells were detected by staining of IE1/2 antigen. The nuclei were counterstained with DAPI. The experiment was repeated two times, and at least 25 foci of each experiment and virus were acquired in the Axio-Observer.Z1 fluorescence microscope with the 10× lens objective.

### Immunocytochemistry

Intracellular localisation studies after transient expression were performed in COS-7 cells that were seeded with 3.3×10^4^ cells per well on glass coverslips in a 24-well plate. The next day, cells were transfected with the respective DNAs by using Lipofectamine LTX (Invitrogen) according to the manufacturer’s protocol. At 24 hours post-transfection, cells were fixed with 4% (v/v) paraformaldehyde (PFA) in PBS for 10 min at 4°C and prepared for localisation analysis by using indirect immunofluorescence staining.

For conditional expression during infection, approx. 5×10^6^ MRC-5 cells were transfected by electroporation with 3 µg of DNA of the inducible expression vector for VPS4A-FLAG (pSVT-VPS4-FLAG) using 260 V, 1050 µF and a 0.4 mm electroporation cuvette. Cells were then recovered from the cuvette in DMEM and seeded into a T25 flask. Next day, transfected cells were seeded on µ-Slide 8- well chamber slides (Ibidi) and infected with different viruses at an MOI of 0.5 to 1. Expression of VPS4A-FLAG was induced by addition of 100 ng/mL doxycycline at day 1 of infection and renewed every 24 hours. For immunofluorescence analysis, cells were fixed after the indicated times with 4% (v/v) PFA in PBS for 10 min at 4°C. Indirect immunofluorescence staining for infected cells as well as transfected cells was performed exactly as described previously (Brock et al., 2013; Read et al., 2019). Confocal images of both transfected and HCMV-infected cells were acquired using the 63× lens objective of the Axio-Observer Z1 fluorescence microscope (Zeiss) equipped with Apotome2. The images were processed with Zen 2.3 software (Zeiss).

### Bimolecular fluorescence complementation (BiFC)

The BiFC assay was performed as previously described (Becker and von Einem, 2013). Therefore, MRC- 5 cells were first transfected by electroporation as described in section *immunocytochemistry* with the DNA of conditional expression vectors expressing either YC-VPS4A-FLAG, YC-VPS4A-FLAG-V13D, or YC- VPS4A-FLAG-V64D fusion proteins. Transfected cells were subsequently infected with TB71-YN at an MOI of 1. Expression was induced by addition of doxycycline (250 ng/mL) at day 2 of infection and renewed every 24 hours. Confocal images of Citrine fluorescence of infected cells were acquired using the 63× lens objective of the Axio-Observer Z1 fluorescence microscope (Zeiss) equipped with Apotome2 after their fixation at 5 dpi and indirect immunofluorescence detection of the FLAG-epitope.

### Immunoprecipitation and Immunoblotting

Co-immunoprecipitation (Co-IP) experiments were performed after 48 hours of transient expression in HEK293FT cells. Briefly, 7×10^5^ cells were seeded per well in a 6-well plate and transfected the next day with 2 µg total DNA. At 48 h post-transfection, cells were washed twice with PBS and lysed for 20 min on ice in lysis buffer (50 mM Tris-HCl, pH 8.0, 150 mM NaCl, 5 mM EDTA, 0.5% NP-40, supplemented with protease inhibitors [Roche]). Centrifugation at 13,000×g for 10 min at 4°C was then used to remove cell debris. One aliquot of the lysate was retained as lysate control and the rest was used for Co-IP experiments. Protein G Dynabeads (Thermo Fisher) were washed with binding buffer (0.1M sodium phosphate buffer, 0.01% Tween20) and then conjugated with FLAG antibody (M2, Sigma) according to the manufacturer’s protocol. For immunoprecipitation, the FLAG antibody-conjugated beads were incubated with the lysate for 1 hour at room temperature. After three washes with lysis buffer, the beads were taken up in 20 µL of 2× SDS sample buffer, incubated at 95°C for 10 min and subjected to SDS-PAGE. The subsequent analysis of the interaction partners was performed by immunoblot analysis as previously described (Schauflinger et al., 2011).

### Ultrastructural analysis

Electron microscopy and quantification of virus morphogenesis stages was performed as previously described (Schauflinger et al., 2011). Briefly, HFFs were cultured on carbon-coated sapphire discs and infected at an MOI of 0.5. At 5 days post infection, the cells were immobilised by high-pressure freezing and freeze substituted, embedded in Epon, and ultrathin sections were mounted on copper grids. The sections were imaged with a JEOL JEM-1400 transmission electron microscope equipped with a CCD camera at an acceleration voltage of 120 kV. The quantification of virus morphogenesis stages was done by counting fully enveloped and budding virus particles in arbitrarily chosen areas of the HCMV cVAC. Micrographs from at least 14 cells for each virus from two independent experiments were used for the quantitative analyses.

### Protein purification

Proteins were expressed in *Escherichia coli* T7 Express *lysY*/*I^q^*cells (New England Biolabs) in 2×TY medium at 37°C to an OD_600_ of 0.8–1.2 before inducing protein expression by addition of 0.4 mM isopropyl β-D-thiogalactopyranoside (IPTG). Expression of GST-tagged pUL71 truncations was induced at 37°C for 3 h, whereas expression of GST-VPS4A MIT domain was induced at 22°C for 16 h. Cells were resuspended at 4°C in GSH lysis buffer (20 mM Tris pH 7.5, 300 mM NaCl, 0.5 mM MgCl_2_, 1.4 µM β-mercaptoethanol, 0.05% TWEEN-20) supplemented with 200–400 U bovine DNase I (Merck) and 200 μL EDTA-free protease inhibitor cocktail (Merck) before lysis using a TS series cell disruptor (Constant Systems) at 24 kpsi. Lysates were cleared by centrifugation (40,000×g, 30 min, 4°C) and incubated with glutathione Sepharose 4B (Cytiva) for 1 h at 4°C before extensive washing (≥ 20 column volumes) with GSH wash buffer (20 mM Tris pH 7.5, 300 mM NaCl, 1 mM DTT) and elution using wash buffer supplemented with 25 mM reduced L-glutathione. GST-tagged pUL71 truncations were subjected to size-exclusion chromatography (SEC) using a Superdex 200 16/600 column (Cytiva) equilibrated in GSH wash buffer. For GST-VPS4A MIT domain, the GST tag was removed by supplementing the pooled SEC fractions containing the protein of interest with 0.5 mM EDTA and incubating overnight at 4°C with 50 µg of GST-tagged human rhinovirus 3C protease. Free GST and uncleaved GST-VPS4A MIT domain were captured using glutathione Sepharose and the cleaved complex was subjected to SEC using an S75 10/300 GL column (Cytiva) equilibrated in 50 mM HEPES pH 8, 100 mM NaCl, 0.5 mM Tris(2-carboxyethyl)phosphine (TCEP).

### Isothermal titration calorimetry

Where necessary, proteins were exchanged into ITC buffer (50 mM HEPES pH 8, 100 mM NaCl, 0.5 mM TCEP) by SEC using Superdex 200 (GST-tagged titrants) or 75 (VPS4 MIT domain) 10/300 GL columns (Cytiva), or by extensive dialysis using tubing with a 3.5 kDa molecular weight cut-off (Pierce). Protein concentrations were estimated from A_280_ using extinction coefficients calculated from the protein sequence (Gasteiger et al., 2005). Peptides encoding residues 166–181 of human CHMP6 or various truncations of HCMV pUL71 were purchased from Genscript at >95% purity and the dry peptides were resuspended in ITC buffer to the desired concentration. ITC experiments were performed at 25°C using an MicroCal PEAQ-ITC automated calorimeter (Malvern Panalytical). Titrants (GST-pUL71, GST-CHMP6 or peptides) were titrated into VPS4A MIT domain using either 19×2 μL or 12×3 μL injections (Table S1). Data were analysed using the MicroCal PEAQ-ITC analysis software (Malvern Panalytical) and fitted using a one-site binding model.

### Molecular dynamics and umbrella sampling

Molecular dynamics simulations were performed using GROMACS version 2020.2 (Abraham et al., 2015). The starting model was a pUL71(300–325):VPS4A MIT domain complex predicted using AlphaFold-Multimer (Evans et al., 2021) version 1.2.1 and amino acid substitutions in pUL71 were incorporated using the mutagenesis wizard in PyMOL, selecting the rotamer with the lowest strain. Models were relaxed to resolve side chain clashes by simulation using the Amber99SB-ILDN force field (Hornak et al., 2006; Lindorff-Larsen et al., 2010). The model was placed in an octahedral box at least 1.0 nm larger than the protein in all dimensions, with periodic boundary conditions, that was explicitly solvated using the TIP3P water model (Jorgensen et al., 1983) and 200 mM NaCl, including additional counter-ions where necessary to neutralise the system charge. Energy minimisation was performed using the steepest descents algorithm with a maximum force tolerance of 500 kJ/mol/nm. Equilibration was performed in two phases with position restraints applied to all non-hydrogen protein atoms. Protein and non-protein atoms were coupled to separate temperature baths maintained at 310 K using the Berendsen weak coupling method (Berendsen et al., 1984). The first phase involved simulation for 100 ps under a constant volume (NVT) ensemble, at the start of which initial velocities were generated using a random seed. Following NVT equilibration the system was equilibrated for 100 ps under a constant pressure (NPT) ensemble, with system pressure maintained at 1.0 bar using the isotropic Berendsen barostat (Berendsen et al., 1984). Production simulations were performed over 10 ns using the V-rescale modified Berendsen thermostat (Bussi et al., 2007) and the isotropic Parrinello-Rahman barostat (Parrinello and Rahman, 1981) in the absence of any atomic restraints. Short-range Coulomb and Van der Waals cut- offs of 1.0 nm were employed, while long-range electrostatics were treated using the Particle-Mesh Ewald (PME) method (Darden et al., 1993; Essmann et al., 1995) with cubic (fourth-order) interpolation. Hydrogen bonds were restrained using the LINear Constraint Solver (LINCS) algorithm (Hess et al., 1997). Dispersion correction was applied to energy and pressure terms to account for truncation of Van der Waals terms.

All steps following energy minimisation were performed five times for each pUL71 mutant so that five random seeds were sampled during velocity generation. All trajectories were corrected for periodic boundary conditions such that both protein molecules remained intact throughout the trajectory. The final frame of the trajectory with lowest mean backbone RMSD for the final 8 ns of the simulation was chosen as the starting model for umbrella sampling. These models were placed in a cuboidal box (5.375 × 20.000 × 6.198 nm) with the vector joining the centres of mass (COMs) of pUL71 and VPS4 aligned with the longest (y) box vector. The box was solvated, ions were added, energy minimisation was performed, and the system was equilibrated as described above. Initial configurations for umbrella sampling were generated using steered molecular dynamics. Non-hydrogen atoms of VPS4A, which served as the reference species, were restrained while the pUL71 peptide was pulled via its COM along the y axis using a spring constant of 1000 kJ/mol/nm^2^ and a pull rate of 0.01 nm/ps, to achieve a total COM separation of approximately 5 nm over 500 ps. The Nosé-Hoover thermostat (Hoover, 1985; Nosé, 1984) was combined with the isotropic Parrinello-Rahman barostat (Parrinello and Rahman, 1981) to ensure that a true NPT ensemble was sampled; all other parameters were as above. From these trajectories, snapshots were taken as the starting configurations for umbrella sampling simulations, with window spacing of 0.1 nm from 0–2.5 nm COM separation and 0.2 nm for > 2.5 nm COM separation. Each of these configurations was equilibrated under an NPT ensemble as described above before data collection. Production simulations were performed for 10 ns using the Nosé-Hoover thermostat (Hoover, 1985; Nosé, 1984) and Parrinello-Rahman barostat (Parrinello and Rahman, 1981), with non-hydrogen atoms of VPS4A subject to position restraints. The COM distance between pUL71 and VPS4 was maintained using a harmonic restraint applied to the pUL71 COM with a spring constant of 1000 kJ/mol/nm^2^. Where particular VPS4A:pUL71 COM separations were not sufficiently sampled, additional umbrella sampling simulations were performed in these regions using starting configurations obtained from the original steered MD trajectories.

Umbrella simulations were analysed using the weighted histogram analysis method (Patey and Valleau, 1973; Torrie and Valleau, 1977, 1974) as implemented in GROMACS (Hub et al., 2010) at 310 K, with potential of mean force (PMF) profiles set to zero at a COM distance of four nanometres. Error analysis was performed using 200 bootstraps, in which complete histograms were considered as independent data points, and random weights were assigned to each histogram (“Bayesian bootstrap”) (Hub et al., 2010).

### Data availability

Structural models and molecular dynamics configuration files and trajectories have been deposited in the University of Cambridge Apollo repository (*URL*).

## Acknowledgements

The authors thank C. Sinzger (Ulm University Hospital), A. Calistri (University of Padova), J. Kudla (University of Muenster), Z. Ruzsics (University Medical Center Freiburg) and W. Britt (University of Alabama Birmingham) for supplying reagents. A Titan V graphics card used for this research was donated by the NVIDIA Corporation. BGB was a Wellcome Trust PhD student. This work was supported by a Sir Henry Dale Fellowship (098406/Z/12/B), jointly funded by the Wellcome Trust and the Royal Society (to SCG). For the purpose of open access, the author has applied a Creative Commons Attribution (CC BY) licence to any Author Accepted Manuscript version arising.

Nuclei are outlined in single channel images and shown (DAPI, blue) in merge. Scale bars = 10 µm. (E) Anti-FLAG immunoprecipitation (IP) from cells co-transfected with VPS4A-FLAG and WT, PPAA or V317D pUL71. Samples were immunoblotted using antibodies as shown. (F) Coomassie-stained SDS-PAGE of GST-tagged WT or PPAA mutant pUL71 C-terminal tail (residues 283–361), VPS4A MIT domain (residues 1–84), or GST-tagged CHMP6 MIM2 motif (residues 168–181) purified following bacterial expression. (G) ITC analysis of the interaction between purified VPS4A MIT domain and WT GST- pUL71(283–361) (*left*), PPAA mutant GST-pUL71(283–361) (*middle*), and GST-CHMP6 MIM2 (*right*). For each, the top graph is baseline-corrected differential power as a function of time and the bottom is the normalised binding curve showing integrated changes in enthalpy (ΔH) as function of molar ratio (syringe:cell component). The corresponding dissociation constant (*K*_D_), number of binding sites (N), and enthalpy change (ΔH) for each representative experiment are shown. All experiments were performed at least twice independently, as detailed in Table 1.

---

**Figure S1.**
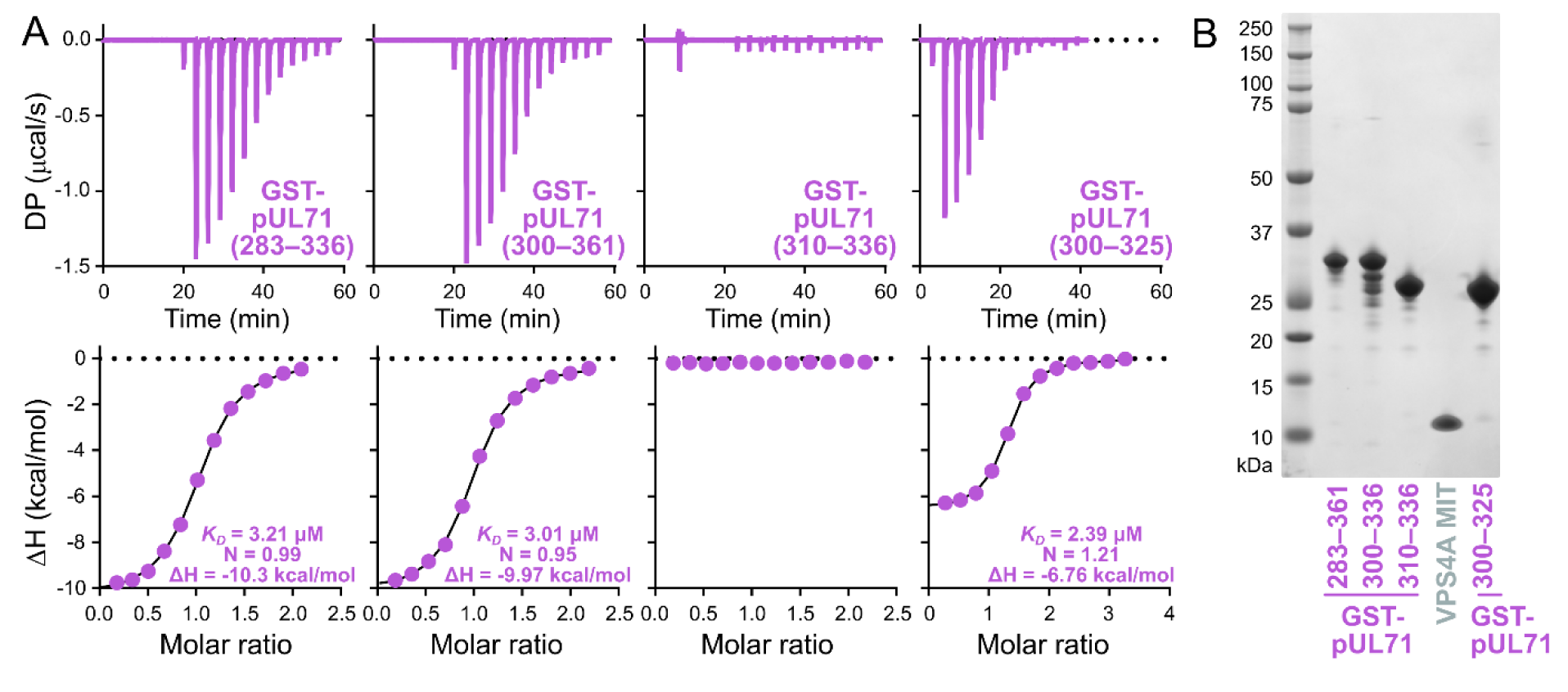
ITC analysis of GST-tagged truncations identifies pUL71 residues 300–325 as necessary and sufficient for binding the VPS4A MIT domain. (A) ITC analysis of the interaction between purified VPS4A MIT domain and GST-tagged truncations of the pUL71 C-terminal tail. (B) Coomassie-stained SDS-PAGE of purified GST-tagged pUL71 C-terminal truncations and VPS4A MIT domain used for ITC analysis.

**Figure S2.**
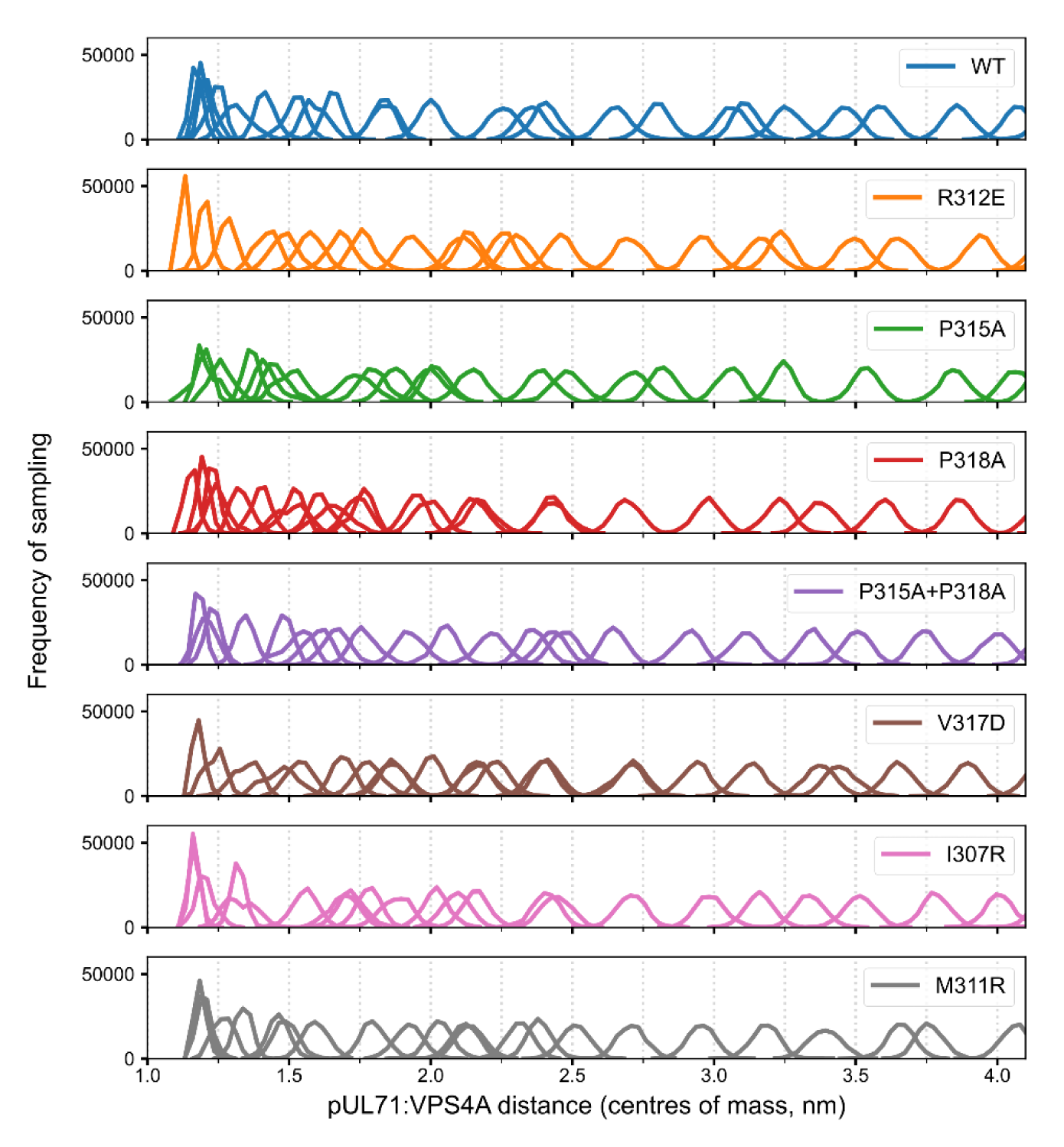
Umbrella sampling of the predicted pUL71(300–325):VPS4A MIT domain interaction. Histograms representing the distribution of pUL71:VPS4A centre-of-mass distances sampled in individual simulations along the reaction coordinate are shown for each pUL71 mutant.

**Figure S3.**
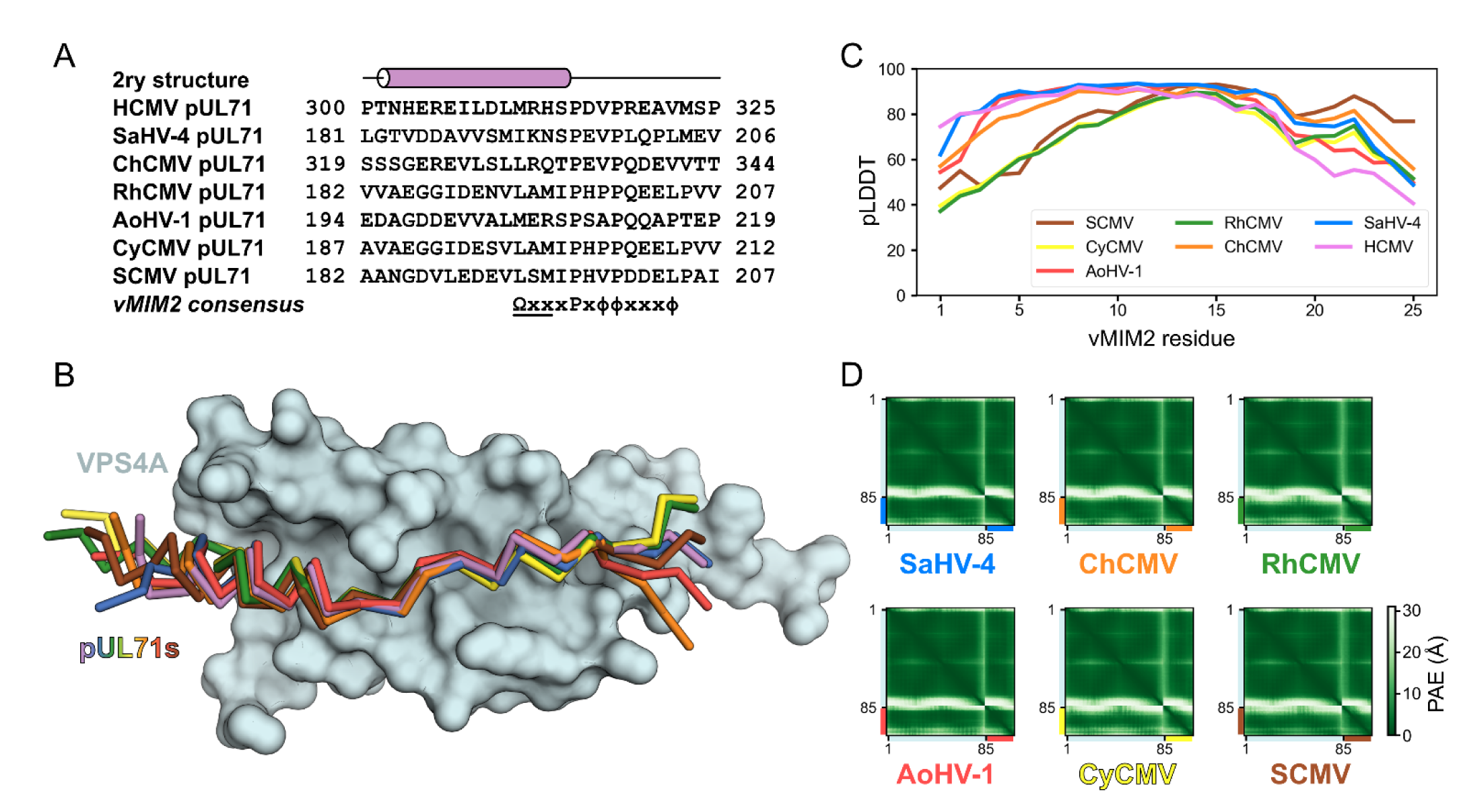
VPS4A binding is conserved across primate cytomegaloviruses. (A) Alignment of vMIM2 sequences of pUL71 homologues from primate cytomegaloviruses: saimiriine betaherpesivirus 4 (SaHV-4), chimpanzee cytomegalovirus (ChCMV), rhesus macaque cytomegalovirus (RhCMV), Aotine betaherpesvirus 1 (AoHV-1), cynomolgus macaque cytomegalovirus (CyCMV), and simian cytomegalovirus (SCMV). The secondary structure of the pUL71 predicted structure is shown above. The pUL71 homologue vMIM2 consensus sequence is shown below, where Ω denotes a large hydrophobic residue, x denotes any residue, ϕ denotes a small hydrophobic residue (including proline), and where the underlined residues are within an α-helix. (B) Superposition of the predicted structure pUL71 vMIM2s (C^α^ traces) from SaHV-4 (blue), ChCMV (orange), RhCMV (green), AoHV-1 (red), CyCMV (yellow) and SCMV (brown) onto the prediction of human cytomegalovirus (HCMV, violet) in complex with human VPS4A MIT domain (cyan molecular surface). Predictions were performed using the VPS4A MIT domain sequence from the cognate host species for each virus, but for clarity only the human VPS4A MIT domain is shown. (C) Per-residue pLDDT scores of residues in the vMIM2s are shown. (D) PAE matrices for predicted pUL71 vMIM2:VPS4 MIT domain complexes.

**Figure S4.**
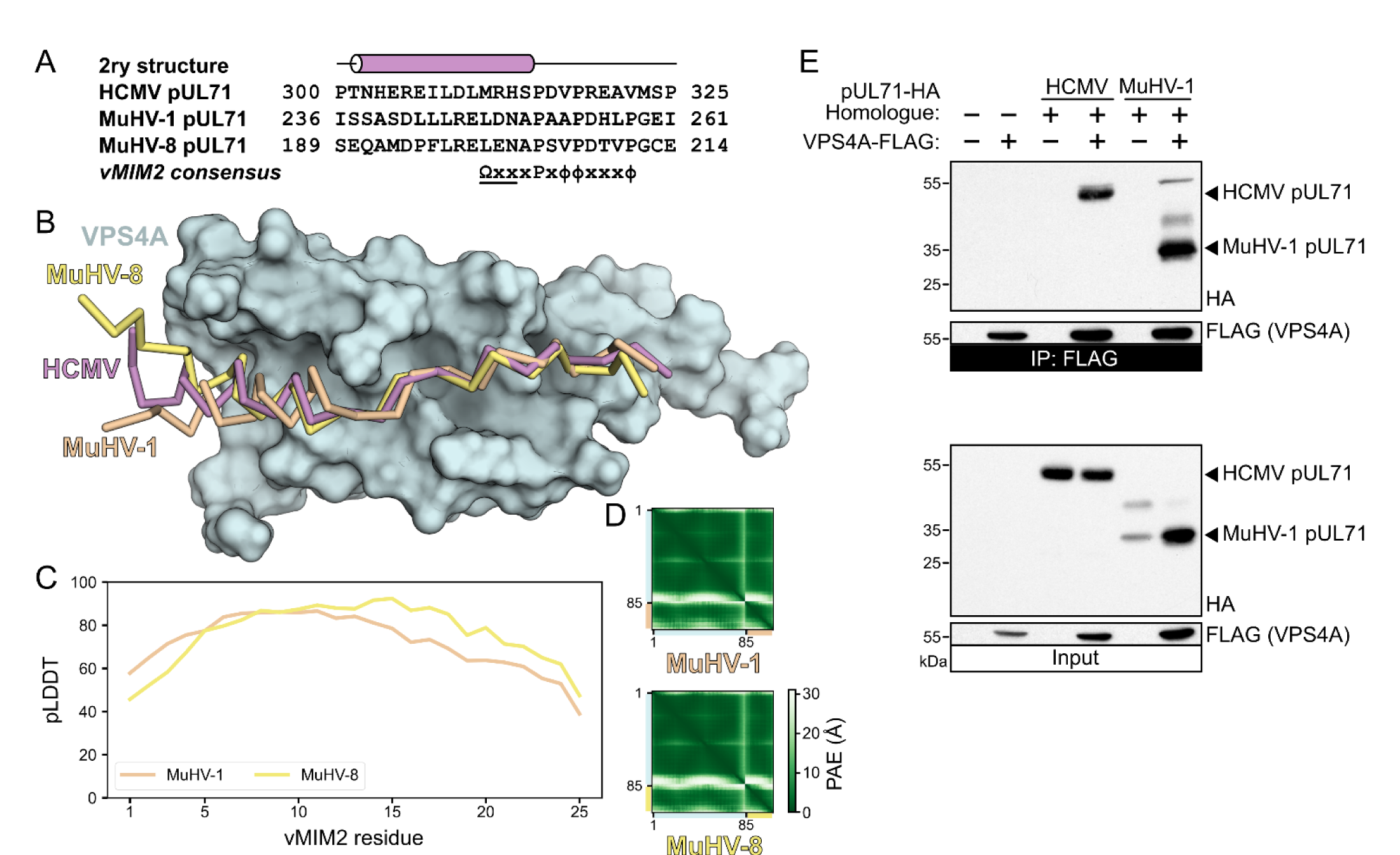
VPS4A binding is conserved across rodent cytomegaloviruses. (A) Alignment of vMIM2 sequences of pUL71 homologues from mouse cytomegalovirus (MuHV-1) and the England isolate of rat cytomegalovirus (MuHV-8). The secondary structure of the pUL71 predicted structure is shown above. The pUL71 homologue vMIM2 consensus sequence is shown below, where Ω denotes a large hydrophobic residue, x denotes any residue, ϕ denotes a small hydrophobic residue (including proline), and where the underlined residues are within an α-helix. (B) Superposition of the predicted structures of pUL71 vMIM2s (C^α^ traces) from MuHV-1 (brown) and MuHV-8 (yellow) onto the prediction of human cytomegalovirus pUL71 (HCMV, violet) in complex with the human VPS4A MIT domain (cyan molecular surface). Predictions were performed using the VPS4A sequence from the cognate host species for each virus, but for clarity only the human VPS4A MIT domain is shown. (C) Per-residue pLDDT scores of residues in the vMIM2s are shown. (D) PAE matrices for predicted pUL71 vMIM2:VPS4 MIT domain complexes. (E) Anti-FLAG immunoprecipitation (IP) from cells co-transfected with human VPS4A-FLAG and HA-tagged pUL71 from HCMV and MuHV-1. Samples were immunoblotted using antibodies as shown.

**Table S1.**
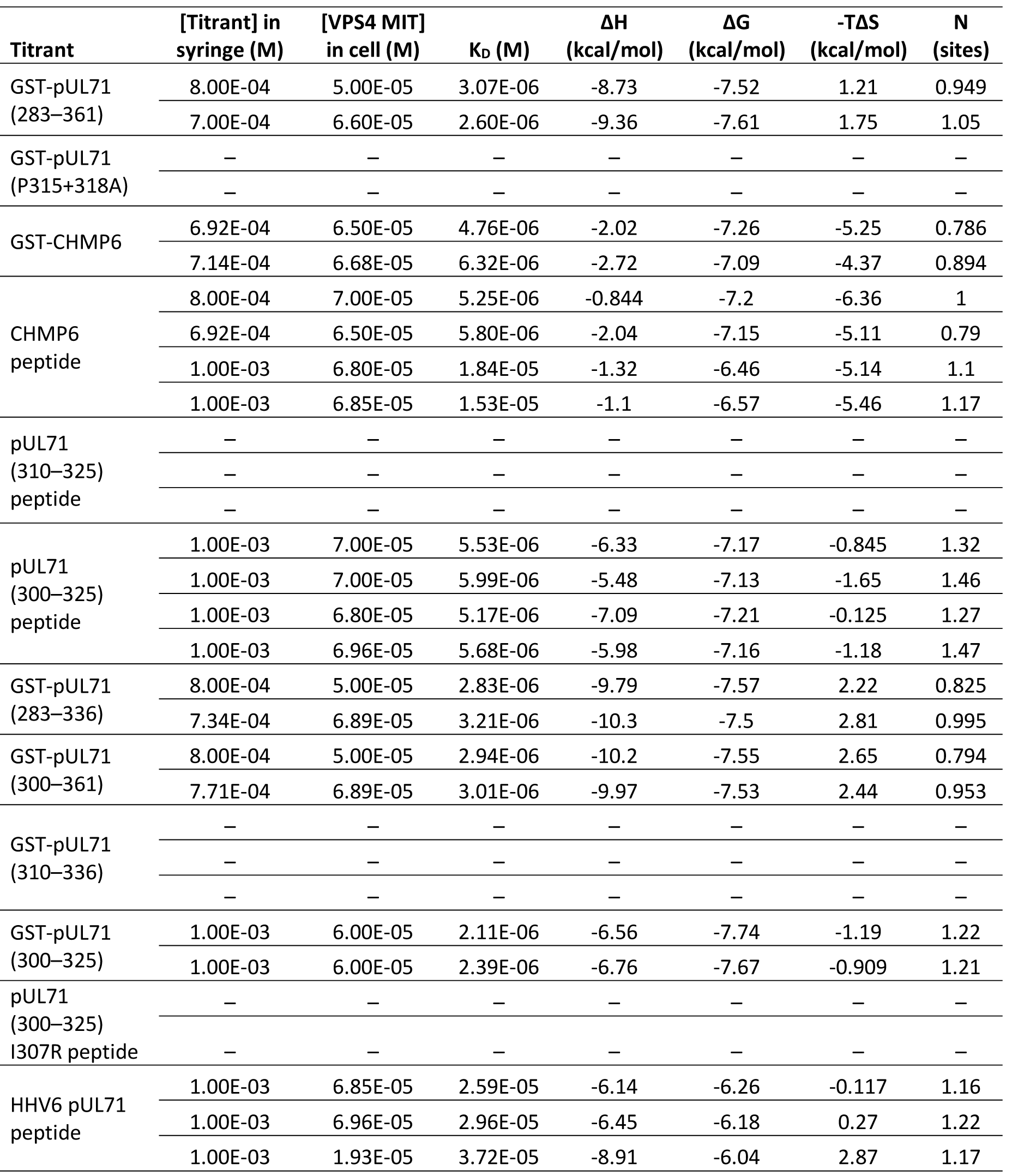
Thermodynamic properties of the interactions with VPS4 MIT domain. As quantitated by isothermal titration calorimetry (ITC). Data for independent experiments are shown. For all, the cell contained human VPS4 MIT domain (residues 1-84) and pUL71 is from HCMV unless stated otherwise. –, no binding detected.

